# ORGaNICs: A Canonical Neural Circuit Computation

**DOI:** 10.1101/506337

**Authors:** David J. Heeger, Wayne E. Mackey

## Abstract

A theory of cortical function is proposed, based on a family of recurrent neural circuits, called ORGaNICs (Oscillatory Recurrent GAted Neural Integrator Circuits). Here, the theory is applied to working memory and motor control. Working memory is a cognitive process for temporarily maintaining and manipulating information. Most empirical neuroscience research on working memory has measured sustained activity during delayed-response tasks, and most models of working memory are designed to explain sustained activity. But this focus on sustained activity (i.e., maintenance) ignores manipulation, and there are a variety of experimental results that are difficult to reconcile with sustained activity. ORGaNICs can be used to explain the complex dynamics of activity, and ORGaNICs can be use to manipulate (as well as maintain) information during a working memory task. The theory provides a means for reading out information from the dynamically varying responses at any point in time, in spite of the complex dynamics. When applied to motor systems, ORGaNICs can be used to convert spatial patterns of premotor activity to temporal profiles of motor control activity: different spatial patterns of pre-motor activity evoke different motor control dynamics. ORGaNICs offer a novel conceptual framework; Rethinking cortical computation in these terms should have widespread implications, motivating a variety of experiments.

## Introduction

Working memory involves much more than simply holding a piece of information online. In cognitive psychology, the idea of working memory includes manipulating online information dynamically in the context of new sensory input. For example, understanding a complex utterance (with multiple phrases) often involves disambiguating the syntax and/or semantics of the beginning of the utterance based on information at the end of the sentence. Doing so necessitates representing and manipulating long-term dependencies, i.e., maintaining a representation of the ambiguous information, and then changing that representation when the ambiguity is resolved.

Most of the empirical neuroscience research on working memory, by contrast, has focused only on maintenance, not manipulation, during delayed-response tasks (1-4). A large body of experimental research has measured sustained activity in prefrontal cortex (PFC) and/or parietal cortex during the delay periods of various such tasks including memory-guided saccades (5-10) and delayed-discrimination and delayed match-to-sample tasks (11-14). Most of the models of working memory, based on neural integrators (see Supplementary Materials, **Figs. S1-S3**, for a primer on neural integrators and neural oscillators), are aimed to explain sustained delay-period activity or to explain well-established behavioral phenomena associated with sustained activity (15-19). There are, however, a variety of experimental results that are difficult to reconcile with sustained activity and neural integrator models. First, some (if not the majority of) neurons either exhibit sequential activity during delay periods such that the activity is handed off from one neuron to the next during a delay period with each individual neuron being active only transiently (20-25), or they exhibit complex dynamics during the delay periods (25-31), not just constant, sustained activity. Second, complex dynamics (including oscillations) are evident also in the synchronous activity (e.g., as measured with local field potentials) of populations of neurons (32, 33). Third, some of the same neurons exhibit activity that is dependent on task demands (34-37). Fourth, some of these neurons appear to contribute to different cognitive processes (controlling attention, decision making, motor control), in addition to working memory, either for different tasks or during different phases of task execution over time (38-42).

Long Short Term Memory units (LSTMs) are machine learning (ML) / artificial intelligence (AI) algorithms that are capable of representing and manipulating long-term dependencies (43), in a manner that is analogous to the concept of working memory in psychology. LSTMs are a class of recurrent neural networks (RNNs). A number of variants of the basic LSTM architecture have been developed and tested for a variety of AI applications including language modeling, neural machine translation, and speech recognition (44-52). (See also: http://colah.github.io/posts/2015-08-Understanding-LSTMs/ and http://karpathy.github.io/2015/05/21/rnn-effectiveness/.) In these and other tasks, the input stimuli contain information across multiple timescales, but the ongoing presentation of stimuli makes it difficult to correctly combine that information over time (53, 54). An LSTM handles this problem by updating its internal state over time with a pair of gates: the update gate selects which part(s) of the current input to process, and the reset gate selectively deletes part(s) of the current output. The gates are computed at each time step from the current inputs and outputs. This enables LSTMs to maintain and manipulate a representation of some of the inputs, until needed, without interference from other inputs that come later in time.

Here, we describe a neurobiological theory of working memory, based on a recurrent neural circuit that we call ORGaNICs (**O**scillatory **R**ecurrent **Ga**ted **N**eural **I**ntegrator **C**ircuit**s**). The theory is an extension of Heeger’s Theory of Cortical Function (55). Because they have the capability of an LSTM, ORGaNICs can solve tasks that are much more sophisticated than the typical delayed-response tasks used in most cognitive psychology and neuroscience experiments. ORGaNICs can exhibit complex dynamics, including sequential activity, but the theory provides a means for reading out information from the dynamically varying responses at any point in time, in spite of the complex dynamics. ORGaNICs can be implemented with a simplified biophysical (equivalent electrical circuit) model of pyramidal cells with separate compartments for the soma, apical dendrite, and basal dendrite. ORGaNICs are also applicable to motor preparation and motor control because ORGaNICs may be used to generate signals with complex dynamics.

A preliminary version of some of this work, along with further details and mathematical derivations was posted on a preprint server (56).

## Results

### ORGaNICs

An example ORGaNICs circuit is depicted in **Fig. 1**. The neural responses of a population of neurons are modeled as dynamical processes that evolve over time. The output responses depend on an input drive (a weighted sum of the responses of a population of input neurons) and a recurrent drive (a recurrent weighted sum their own responses). The time-varying output responses are represented by a vector **y** = (*y*_*1*_, *y*_*2*_,…, *y*_*j*_,…, *y*_*N*_) where the subscript *j* indexes different neurons in the population. (We use boldface lowercase letters to represent vectors and boldface uppercase to denote matrices.) The timevarying inputs are represented by another vector **x** = (*x*_*1*_, *x*_*2*_,…, *x*_*j*_,…, *x*_*M*_). The output responses are also modulated by two populations of time-varying modulators, recurrent modulators **a** and input modulators **b**. (We use the term “modulator” to mean a multiplicative computation regardless of whether or not it is implemented with neuromodulators) Each of these modulators depends on the inputs and outputs. So there are two nested recurrent circuits. 1) Recurrent drive: the output responses depend on the recurrent drive, which depends on a weighted sum of their own responses. 2) Multiplicative modulators: the output responses are modulated (multiplicatively) by the responses of two other populations of neurons (the modulators), which also depend on the output responses.

**Figure 1.**
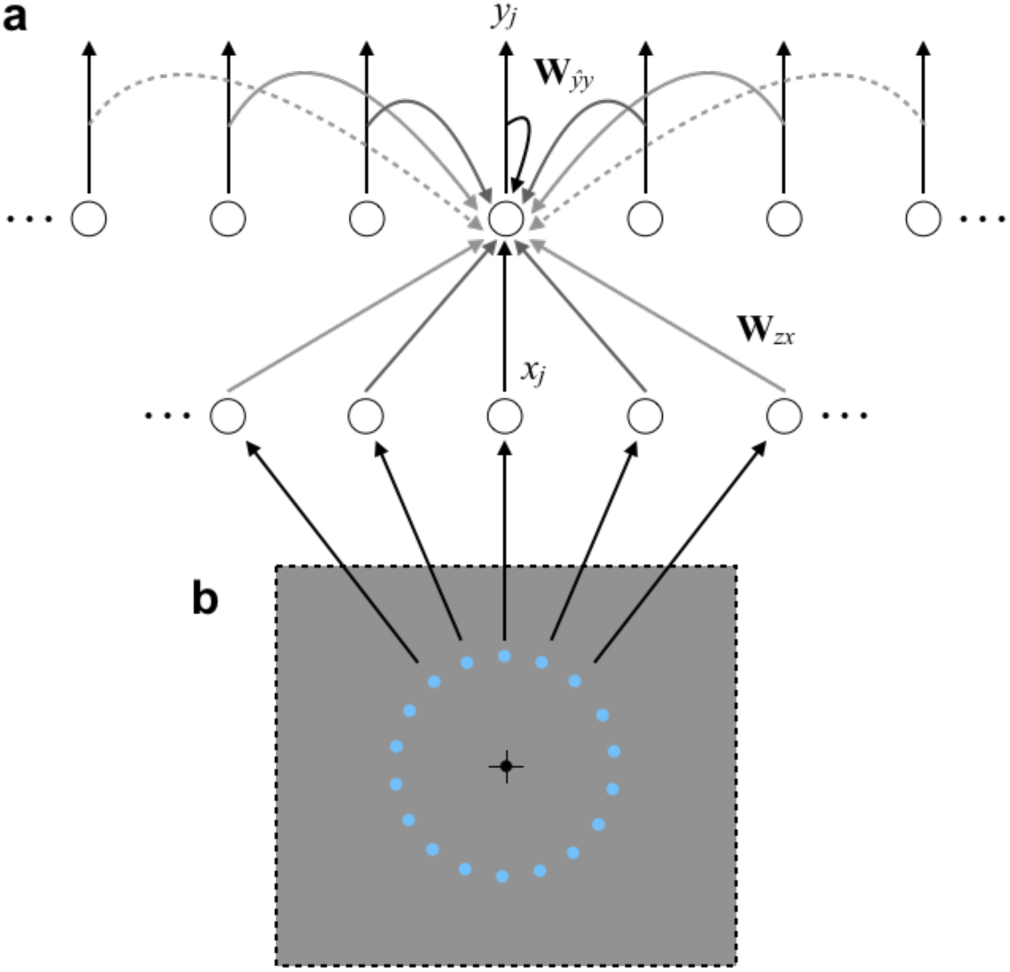
ORGaNICs architecture. **a.** Diagram of connections in an example ORGaNIC. Solid lines/curves are excitatory (positive weights) and dashed curves are inhibitory (negative weights). Gray scale represents strength of connections (weight magnitude). Only a few of the feedforward and recurrent connections are shown to minimize clutter. Modulatory connections not shown. **b.** Oculomotor delayed response task. Black crosshairs, fixation point and eye position at the beginning of each trial. Blue circles, possible target locations, each of which evokes an input.

Specifically, neural responses are modeled by the following dynamical systems equation:

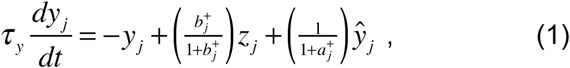

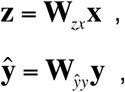

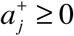 and 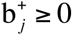

The variables (**y**, **ŷ**, **x**, **z**, **a**, and **b**) are each functions of time, e.g., **y**(*t*), but we drop the explicit dependence on *t* to simplify the notation. The responses **y** depend on an input drive **z**, which is computed as a weighted sum of inputs **x**. The encoding weight matrix (also called the embedding matrix) **W**_*zx*_ is an *N*x*M* matrix of weights where *N* is the number of neurons in the circuit and *M* is the number of inputs to the circuit. The rows of **W**_*zx*_ are the receptive fields of the neurons. The responses **y** also depend on a recurrent drive **ŷ**, which is computed as a weighted sum of the responses **y**. The recurrent weight matrix **W**_*ŷy*_ is an *N*x*N* matrix. If **W**_*ŷy*_ is the identity matrix, then each neuron receives a recurrent excitatory connection from itself. If **W**_*ŷy*_ has a diagonal structure, then each neuron receives recurrent connections from itself and its neighbors. This could, for example, be a center-surround architecture in which the closest recurrent connections are excitatory and the more distant ones are inhibitory (e.g., as depicted in **Fig. 1**). The recurrent drive and input drive are modulated, respectively, by two other populations of neurons: the recurrent modulators **a** and the input modulators **b**. The superscript + is a rectifying output nonlinearity. Halfwave rectification is the simplest form of this rectifying nonlinearity, but other output nonlinearities could be substituted, e.g., sigmoid, exponentiation, half-squaring (57), normalization (58, 59), etc. The value of *τ*_*y*_ is the intrinsic time-constant of the neurons. Finally, the output responses are multiplied by a readout matrix, **r = W**_*ry*_ **y**, where **r** is the readout over time and **W**_*ry*_ is the readout matrix.

The modulators **a** and **b** are themselves modeled as dynamical systems that depend on weighted sums of the inputs and outputs, for example:

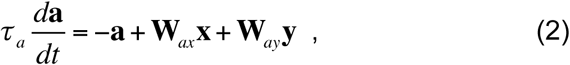

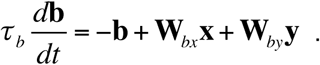

The values of *τ*_*a*_ and *τ*_*b*_ are the intrinsic time-constant of the neurons.

The following subsections illustrate some of the different operating regimes of ORGaNICs. The weights in the various weight matrices (**W**_*zx*_, **W**_*ŷy*_, **W**_*ry*_, **W**_*ax*_, **W**_*ay*_, **W**_*bx*_, **W**_*by*_) are presumed to be learned, depending on task demands. But the elements of these weight matrices were pre-specified (not learned) for each of the simulations in this paper. The time-varying values of the modulators, **a** and **b**, determine the state of the circuit by controlling the recurrent gain and effective time-constant. If both *a*_*j*_ and *b*_*j*_ are large then the response time-course is dominated by the input drive, so the response time-course exhibits a short effective time constant. If both *a*_*j*_ and *b*_*j*_ are small (near 0) then the responses are dominated by the recurrent drive, so the response time-course exhibits a long effective time constant. If *a*_*j*_ is large and *b*_*j*_ is small then the recurrent drive is shut down (like the reset gate in an LSTM). A leaky neural integrator corresponds to a special case in which *a*_*j*_ = *b*_*j*_ is constant over time (see Supplementary Materials for a primer on neural integrators). For some choices of the recurrent weight matrix, the circuit can exhibit sustained activity. For other choices of the recurrent weight matrix, the circuit can exhibit either stable, ongoing oscillations or sequential activity.

### Sustained activity

A large body of experimental research has measured sustained activity in PFC and/or parietal cortex during the delay periods of of delay-response tasks, and most of the models of working memory are aimed to explain sustained delay-period activity (see Introduction for references).

We, likewise, used ORGaNICs to simulate sustained delay-period activity during a memory-guided saccade task (**Fig. 2**), using the circuit depicted in **Fig. 1a**. In this task, a target is flashed briefly while a subject is fixating the center of a screen (**Fig. 1b**). After a delay period of several seconds, the fixation point disappears cueing the subject to make an eye movement to the remembered location of the target. Each neuron in the simulation responded selectively to target location, each with a different preferred polar angle (i.e., saccade direction) in the visual field (**Figs. 1b** and **2a**), all with the same preferred radial position (i.e., saccade amplitude). We ignored saccade amplitude for this simulation, but it would be straightforward to replicate the circuit for each of several saccade amplitudes. The input drive **z** to each neuron, consequently, depended on target position and the time course of the target presentation (**Figs. 2d** and **2e**). The recurrent weights **W**_*ŷy*_ were chosen to have a center-surround architecture; each row of **W**_*ŷy*_ had a large positive value along the diagonal (self-excitation), flanked by smaller positive values, and surrounded by small negative values (**Fig. 2b**). All neurons in the circuit shared the same pair of modulators (*a*_*j*_ = *a* and *b*_*j*_ = *b*), i.e., all of the neurons had the same state at any given point in time. The input to the circuit also comprised the time courses of two cues, one of which indicated the beginning of the trial (at time 0 ms) and the other of which indicated the end of the delay period (at time 3000 ms). The response time-courses of the modulators followed the two cues (**Figs. 2f**), by setting appropriate values in the weight matrices **W**_*ax*_ and **W**_*bx*_. The modulators did not depend on the output responses, i.e., **W**_*ay*_=**0**, **W**_*by*_=**0**.

**Figure 2.**
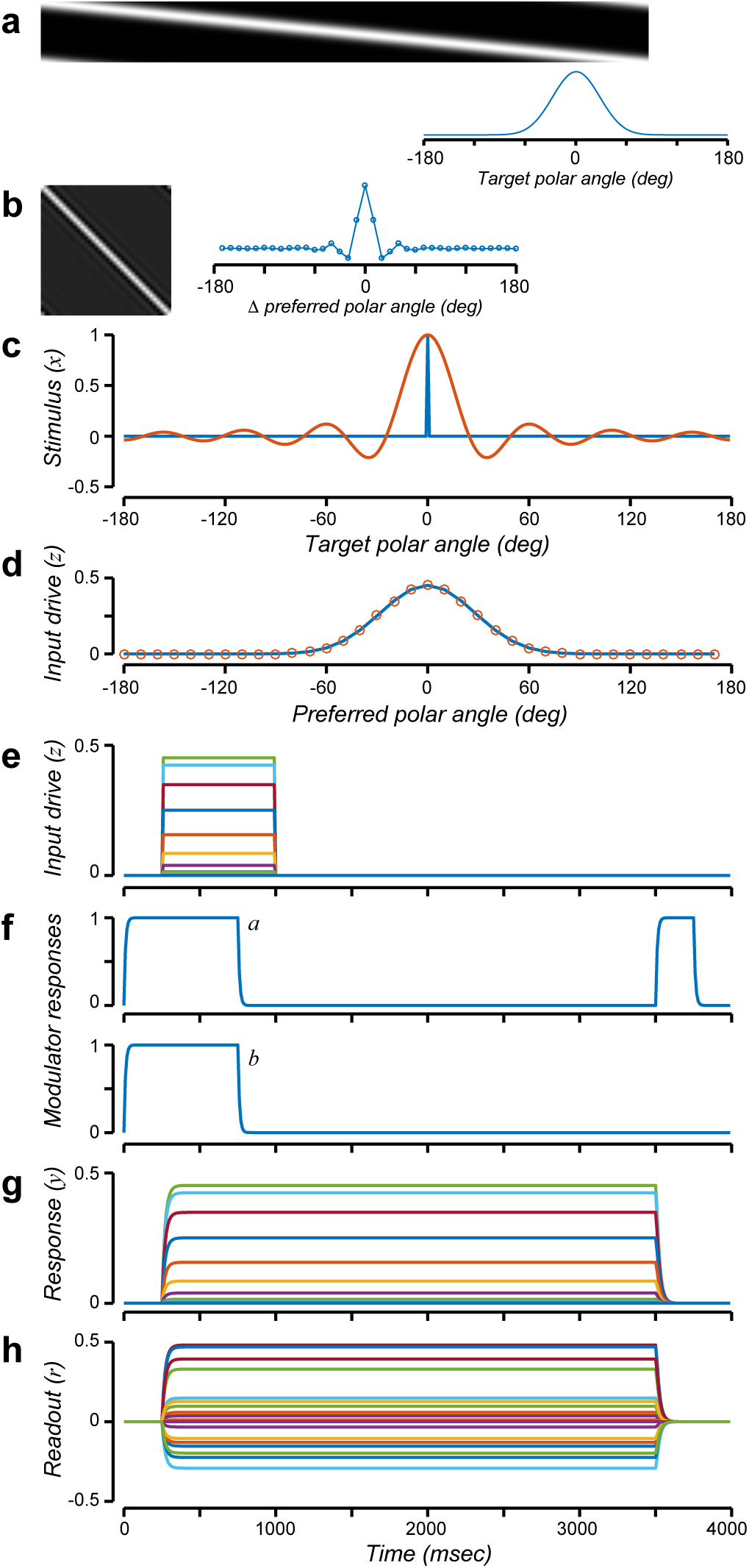
Sustained activity. **a.** Encoding matrix (**W**_*zx*_), each row of which corresponds to a neuron’s receptive field. Graph, encoding weights corresponding to the middle row of the matrix. **b.** Recurrent weight matrix (**W**_*ŷy*_), each row of which corresponds to the recurrent synaptic weights from other neurons in the population. Graph, recurrent weights corresponding to the middle row of the matrix. **c.** Input stimulus and reconstructed stimulus. Blue, input stimulus (**x**) corresponding to target position. Orange, reconstructed stimulus, computed as a weighted sum of the reconstructed input drive (panel d). **d.** Input drive and reconstructed input drive. Blue, input drive (**z**) to each neuron as a function of that neuron’s preferred target location. Orange, reconstructed input drive, computed as a weighted sum of the readout (panel h). **e.** Input drive (**z**) over time. Each color corresponds to a different neuron. **f.** Modulator responses. Top row, *a*. Bottom row, *b*. **g.** Output responses (**y**). Each color corresponds to a different neuron. **h.** Readout (**r**). Each color corresponds to a different component of the readout.

This circuit was capable of maintaining a representation of target location during the delay period with sustained activity (**Fig. 2g**). The responses followed the input drive at the beginning of the simulated trial (compare **Figs. 2e** and **2g**) because *a* and *b* were large (=1, corresponding to a short effective time constant). The values of *a* and *b* then switched to be small (=0, corresponding to a long effective time constant) before the target was extinguished, so the output responses exhibited sustained activity (**Figs. 2g**). Finally, the value of *a* was then switched back to be large (=1, corresponding to a small recurrent gain) at the end of the trial, causing the output responses to be extinguished.

The input drive and target location were reconstructed from the responses, at any time during the delay period (**Figs. 2c** and **2d**). To do so, the responses were first multiplied by a readout matrix, **r = W**_*ry*_ **y**, where **r** is the readout over time (**Fig. 2h**) and **W**_*ry*_ is the readout matrix (computed as described below). The readout, at any time point, was then multiplied by a decoding (or reconstruction) matrix (see Supplementary Material). The result was a perfect reconstruction of the input drive (**Fig. 2d**, orange), and an approximate reconstruction of the input stimulus (**Fig. 2c**, orange) with a peak at the target location. The reconstruction of the input stimulus was imperfect because the receptive fields were broadly tuned for polar angle. Regardless, we do not mean to imply that the brain attempts to reconstruct the stimulus from the responses. The reconstruction merely demonstrates that the responses **y** and the readout **r** implicitly represent the target location.

The dynamics of the responses, during the delay period, depended on the eigenvalues and eigenvectors of the recurrent weight matrix **W**_*ŷy*_. The recurrent weight matrix (**Fig. 2b**) was a symmetric 36×36 matrix (*N*=36 was the number of neurons in the circuit, i.e., each of **y** and **z** were 36-dimensional vectors). Nineteen of the eigenvalues were equal to 1, and the others had values between 0 and 1. The weight matrix was in fact scaled so that the largest eigenvalues were equal to 1. The corresponding eigenvectors defined an orthonormal coordinate system (or basis) for the responses. The responses during the delay period (when *a*=0 and *b*=0) were determined entirely by the projection of the initial values (the responses at the very beginning of the delay period) onto the eigenvectors. Eigenvectors with corresponding eigenvalues equal to 1 were sustained throughout the delay period. Those with eigenvalues less than 1 decayed to zero (smaller eigenvalues decayed more quickly). Those with eigenvalues greater than 1 would have been unstable, growing without bound (which is why the weight matrix was scaled so that the largest eigenvalues = 1).

The readout matrix **W**_*ry*_ = **V**^*t*^ was a 19×36 matrix, where the rows of **V**^*t*^ were computed from the eigenvectors of the recurrent weight matrix **W**_*ŷy*_. Specifically, **V** was an orthonormal basis for the 19-dimensional subspace spanned by the eigenvectors of **W**_*ŷy*_ with corresponding eigenvalues = 1.

The encoding matrix **W**_*zx*_ was a 36×360 matrix (*M*=360 was the number of polar angle samples in the input stimulus). The receptive fields (i.e., the rows of the encoding weight matrix **W**_*zx*_) were designed based on the same eigenvectors. Doing so guaranteed that the input drive was reconstructed perfectly from the responses at any time during the delay period (**Fig. 2d**; see Supplementary Material for derivation).

The example circuit depicted in **Figs. 1** and **2** had a representational dimensionality *d*=19, because the recurrent weight matrix had 19 eigenvalues = 1. The neural activity in this circuit was a 19-dimensional continuous attractor during the delay period. It could, in principle, maintain up to 19 values corresponding to 19 targets, although fewer can be recovered in practice because the receptive fields are broadly tuned.

### Oscillatory activity

There are a variety of experimental results that are difficult to reconcile with sustained activity and neural integrator models. Many neurons either exhibit sequential activity during delay periods such that the activity is handed off from one neuron to the next during a delay period with each individual neuron being active only transiently, or they exhibit complex dynamics during the delay periods, not just constant, sustained activity (see Introduction for references).

A variant of the same circuit was used to generate delay-period activity with complex dynamics (**Fig. 3**), not simply sustained activity, and the same theoretical framework was used to analyze it. The key idea is that the recurrent weights and the output responses may be complex-valued. The complex-number notation is just a notational convenience (see Supplementary Materials). The complex-valued responses may be represented by pairs of neurons, and the complex-valued weights in the recurrent weight matrix may be represented by pairs of synaptic weights (with each pair of synaptic weights shared by each pair of neurons to perform complexnumber multiplication).

**Figure 3.**
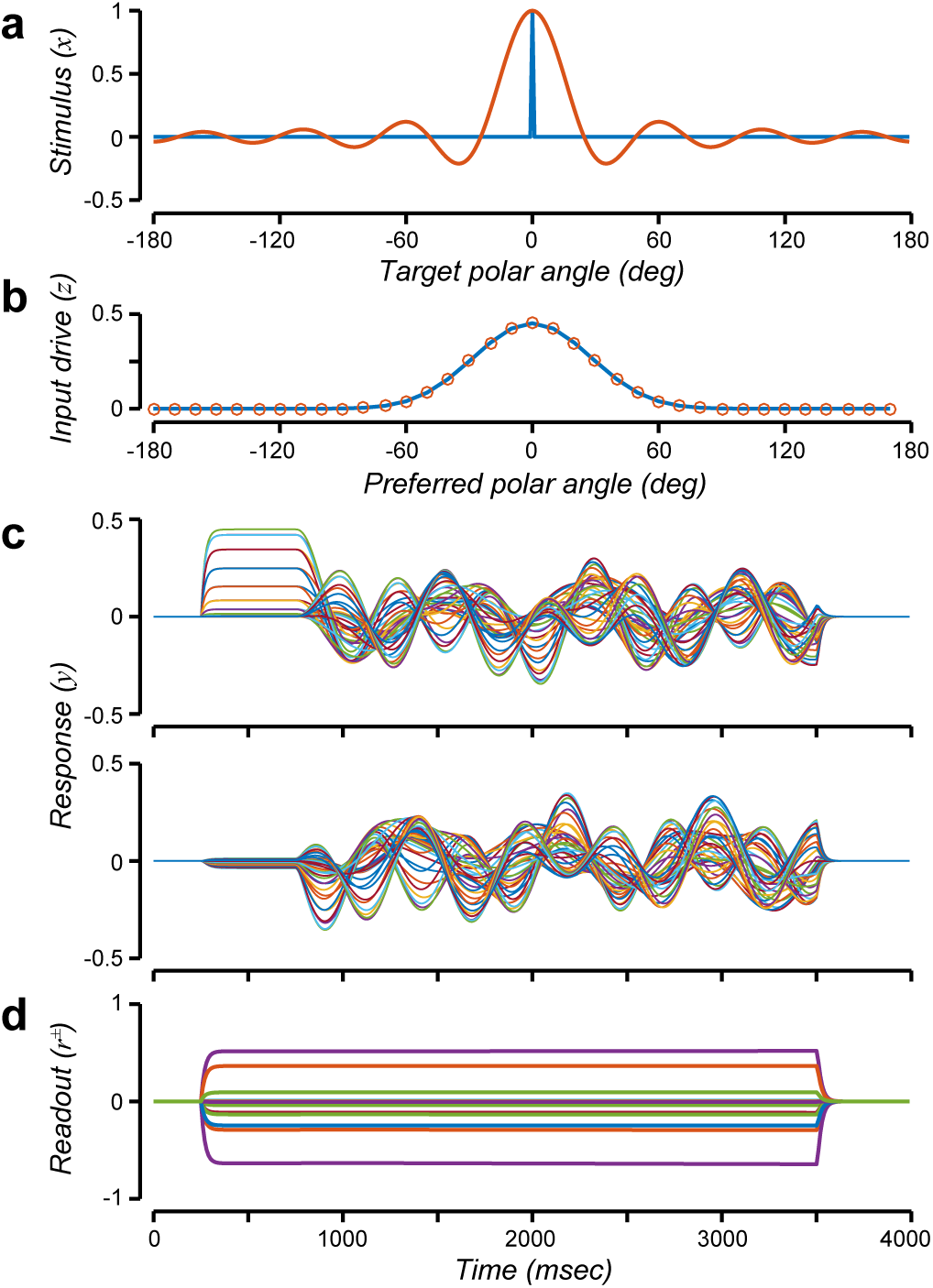
Oscillatory activity. **a.** Input stimulus and reconstructed stimulus. Blue, input stimulus (**x**) corresponding to target position. Orange, reconstructed stimulus, computed as a weighted sum of the reconstructed input drive (panel b). **b.** Input drive and reconstructed input drive. Blue, input drive (**z**) to each neuron as a function of that neuron’s preferred target location. Orange, reconstructed input drive, computed as a weighted sum of the readout (panel d) at a randomly chosen time point during the delay period. **c.** Output responses (**y**). Top row, real part of the responses. Bottom row, imaginary part of the responses. Each color corresponds to a different neuron. **d.** Readout (**r**^±^). Each color corresponds to a different component of the readout.

In this example circuit, there were again 36 neurons with the same encoding matrix **W**_*zx*_ as in the preceding (sustained activity) example. The modulators were also the same as in the preceding example. The real part of the recurrent weight matrix **W**_*ŷy*_ was the same as that in the preceding example (**Fig. 2a**), such that 19 eigenvalues had real parts equal to 1, and the real parts of the other eigenvalues were less than 1. But the imaginary part of the recurrent weight matrix was different, with random values (between 0 and 4π*τ*_*y*_) for the imaginary parts of all 36 eigenvalues, corresponding to oscillation frequencies between 0 and 2 Hz (although other frequencies or combinations of frequencies could be used instead). Consequently, the activity exhibited complex dynamics (sums of oscillations) that resembled the phenomenology of delay-period activity (26-31). The circuit was a 19-dimensional (complex-valued) continuous attractor during the delay period because the recurrent weight matrix was constructed to have 19 eigenvalues with real parts equal to 1.

In spite of the complex response dynamics (**Fig. 3c**), the readout was again (as it was for the sustained activity circuit) constant over time during the delay period (**Fig. 3d**), the input drive was reconstructed perfectly from the responses at any time during the delay period (**Fig. 3b**), and target location was reconstructed approximately (**Fig. 3a**). The readout **r**^±^ for this example circuit was more complicated than that for the sustained activity circuit; it depended both a weighted sum of the responses and an estimate of the sign. The readout matrix (the weighted sum) was computed as a unitary basis for the subspace spanned by the eigenvectors of **W**_*ŷy*_ with corresponding eigenvalues that had real parts = 1. The sign was computed from the output responses (see Supplementary Material).

### Sequential activity

Another variant of the same circuit was used to generate sequential activity (**Fig. 4**). In this example circuit, there were again 36 neurons with the same encoding matrix **W**_*zx*_ as in the preceding examples. The modulators were also the same as in the preceding examples. The recurrent weight matrix was real-valued but asymmetric (**Fig. 4a**, see Supplementary Material). Because of the asymmetry, the eigenvectors and eigenvalues of the recurrent weight matrix were complex-valued, and the output responses exhibited oscillatory dynamics (**Fig. 4b**). In particular, the recurrent weight matrix was designed so that the activity was handed off from one neuron to the next during the delay period, analogous to a synfire chain (60-64), but with continuous activity over time.

**Figure 4.**
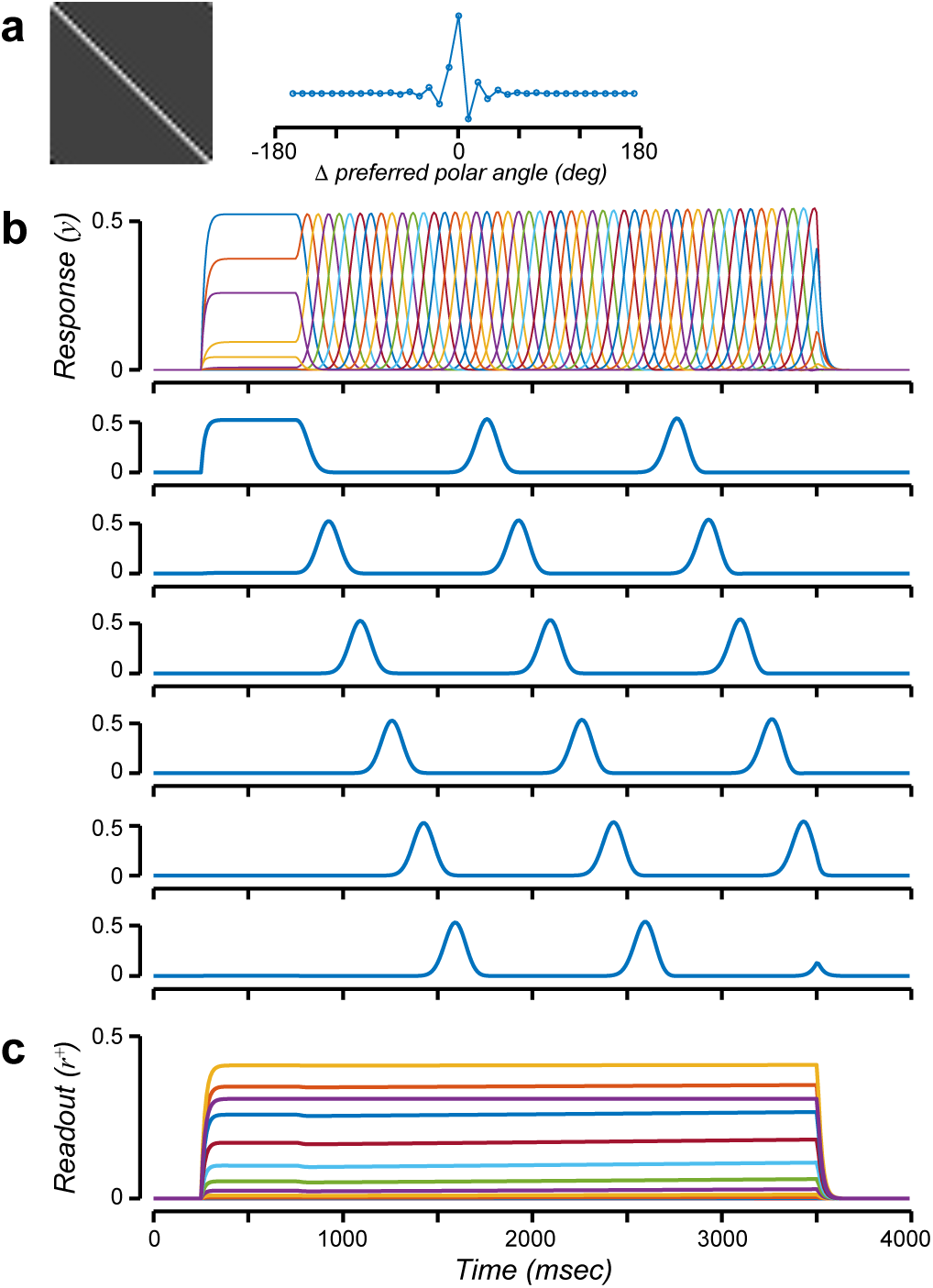
Sequential activity. **a.** Recurrent weight matrix (**W**_*ŷy*_). Graph, recurrent weights corresponding to the middle row of the matrix. **b.** Output responses (**y**). Each color corresponds to a different neuron. Successive rows, responses of a few example neurons. **c.** Readout (**r**^+^). Each color corresponds to a different component of the readout.

In spite of the complex dynamics, the readout was constant over time (**Fig. 4c**). The readout matrix was again, as for the preceding examples, computed as a unitary basis for the subspace spanned by the eigenvectors of **W**_*ŷy*_ with corresponding ei-genvalues that had real parts = 1. But the readout was computed as **r^+^ = |W**_*ry*_ **y|**, i.e., the modulus (square-root of the sum of squares of real and imaginary parts) of a weighted sum of the responses. Consequently, this circuit was capable of maintaining some (but not all) information about the input during the delay period. Unlike the preceding examples, it was not possible to reconstruct the input drive from the readout at arbitrary points in time during the delay period. A linear reconstruction (like that used for the preceding examples) generated a shifted copy of the input drive, that shifted over time like a traveling wave (see Supplementary Material, **Fig. S4**). That is, the information maintained during the delay period was sufficient for discriminating some inputs (e.g., two targets with different intensities or two pairs of targets with different spacings), but incapable of discriminating between other inputs (e.g., a single target of the same intensity presented at two different polar angle locations).

### Motor preparation and motor control

ORGaNICs are also capable of generating signals, like those needed to execute a complex sequence of movements (e.g., speech, bird song, tennis serve, bowling, skiing moguls, backside double McTwist 1260 on a snowboard out of the halfpipe). Some actions are ballistic (open loop), meaning that they are executed with no sensory feedback during the movement. Others are closed loop, meaning that the movements are adjusted on the fly based on sensory feedback. ORGaNICs evoke patterns of activity over time that might underlie the execution of both open- and closed-loop movements.

An example of open-loop control (**Fig. 5**) was implemented using the sequential activity circuit described above, but with a different readout. The encoding matrix and the recurrent matrix were identical to those in the sequential activity circuit. The modulators were also the same as in the preceding examples. The readout was different, simply summing across the components, *r***^Σ^ =Σ**Re(**W**_*ry*_ **y**). Different spatial patterns of inputs led to different temporal dynamics of the responses. When the input was chosen to drive a particular eigenvector (i.e., because the input drive was orthogonal to the other eigenvectors), then the readout during the period of motor execution (same as the delay period in the preceding example circuits) was a 1 Hz sinusoid (**Fig. 5a**). When the input was chosen to drive another eigenvector, then the readout was an 8 Hz sinusoid (**Fig. 5c**). A linear sum of these inputs evoked a linear sum of the readouts (**Fig. 5d**).

**Figure 5.**
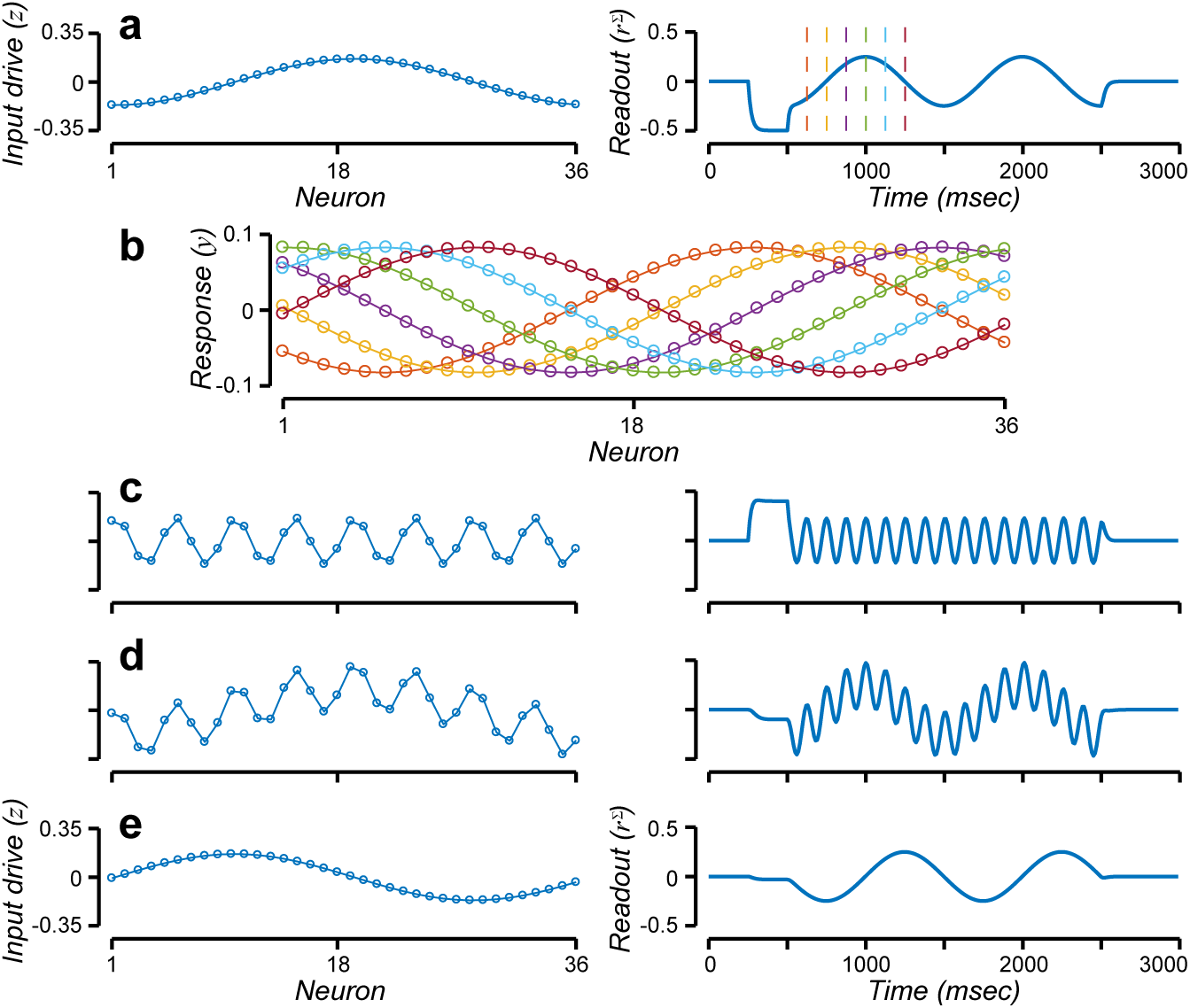
Motor preparation and motor control. **a.** Input drive and readout corresponding to input (with amplitude = 1) that drives only the 1 Hz component of the recurrent weight matrix. Left panel, input drive (**z**), spatial pattern activity across the 36 neurons during the premotor time period (250-500 ms). Right panel, readout (**r^Σ^**) over time. Vertical lines, times corresponding to curves in panel b. **b.** Responses exhibit an oscillating traveling wave of activity. Different colors correspond to different time points, indicated in panel a. **c.** Input drive and readout corresponding to the 8 Hz component of the recurrent weight matrix. Same format as panel a. **d.** Summing the inputs from panels a and b evokes the sum of the responses. **e.** Input drive from panel a is shifted in space, generating a readout that is shifted in time.

How are these temporal profiles of activity generated? Each eigenvector of the recurrent weight matrix is associated with a basis function, a pattern of activity across the population of neurons and over time. Each basis function is a complex exponential (i.e., comprising sine and cosine), the frequency of which is specified by the imaginary part of the corresponding eigenvalue:

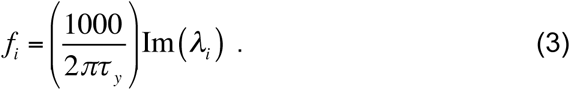

The value of *λ*_*i*_ is the imaginary part of the *i*th eigenvalue of the recurrent weight matrix, and *f*_*i*_ is the corresponding oscillation frequency (in Hz). The factor of 1000 is needed because the time constant *τ*_*y*_ is presumed to be specified in msec but the oscillation frequency is specified in Hz (cycles/sec). The responses exhibit an oscillating traveling wave (**Fig. 5b**); the response of any individual neuron oscillates over time and the entire pattern of activity across the population of neurons shifts over time (**Fig. 5b**, orange - yellow - purple - green - cyan - red). For inputs corresponding to different eigenvectors, the responses oscillate at correspondingly different frequencies (**Fig. 5c**). The frequencies of the various components corresponding to each of the eigenvalues, for this particular recurrent weight matrix, included a number of other frequencies in addition to the 1 and 8 Hz components shown in the figure. Motor control signals with any arbitrary phase, for each of the frequency components, can be generated by shifting the input drive (**Fig. 5e**). That way, all combinations of frequencies and phases can be generated just by changing the spatial pattern of premotor activity, with a fixed, linear readout. This dovetails with experimental evidence demonstrating that the function of motor preparation is to set the initial conditions that generate the desired movement (65-67), and that complex movements are based on a library of motor primitives (68, 69).

The readout for open-loop control is, in general, a linear sum of the responses *r***^Σ^**. The readout matrix for short-term memory, in some of the preceding examples, comprised eigenvectors of the recurrent weight matrix to ensure that the input was recovered at any time during the delay period. But recovering the input is not the goal for open-loop control. Rather, a sum of the (co-)sinusoidal basis functions was used to generate motor control signals for ballistic (open loop) movements.

ORGaNICs are also capable of generating more complicated control signals. The basis functions are damped oscillators when the modulators are greater than 0 but equal to one another (*a* **=** *b*) and constant over time, and when the input is constant over time. If the input is varying over time, then the responses depend on a linear combination of the inputs and the basis functions, and the responses may be used for closed-loop control. If the modulators are also time varying, and different for each neuron, then the responses may exhibit a wide range of dynamics, with the capability (by analogy with LSTMs) of solving relatively sophisticated tasks (see Introduction for references).

### Manipulation: spatial updating

Many tasks require manipulation of information (i.e., working memory) in addition to maintenance of information (i.e., short-term memory). The computational framework of ORGaNICs (by analogy with LSTMs) uses the modulators **a** and **b** to encode a dynamically-changing state that depends on the current inputs and past context. The modulators can depend on the current inputs and outputs, and the current outputs can depend on past inputs and outputs, thereby encoding a state. The modulators can be controlled separately for each neuron so that each neuron can have a different state (different values for *a*_*j*_ and *b*_*j*_) at each instant in time. In the example that follows, however, all the neurons in the circuit shared the same state, but that state changed over time to enable manipulation (via gated integration and reset) in addition to maintenance.

A simulation of the double-step saccade task illustrates how ORGaNICs can both maintain and manipulate information over time (**Fig. 6**). In this task, two targets are shown while a subject is fixating the center of a screen (**Fig. 6a**, upper panel). A pair of eye movements are then made in sequence to each of the two targets. Eye movements are represented in the brain using retinotopic, i.e., eye-centered coordinates (**Fig. 6a**, upper panel, red lines). Consequently, after making the first eye movement, the plan for the second eye movement must be updated (**Fig. 6a**, lower panel, solid red line copied from upper panel no longer points to the second target). This is done by combining a representation of the target location with a copy of the neural signals that control the eye muscles (i.e., corollary discharge) to update the planned eye movement (**Fig. 6a**, lower panel, dashed red line).

**Figure 6.**
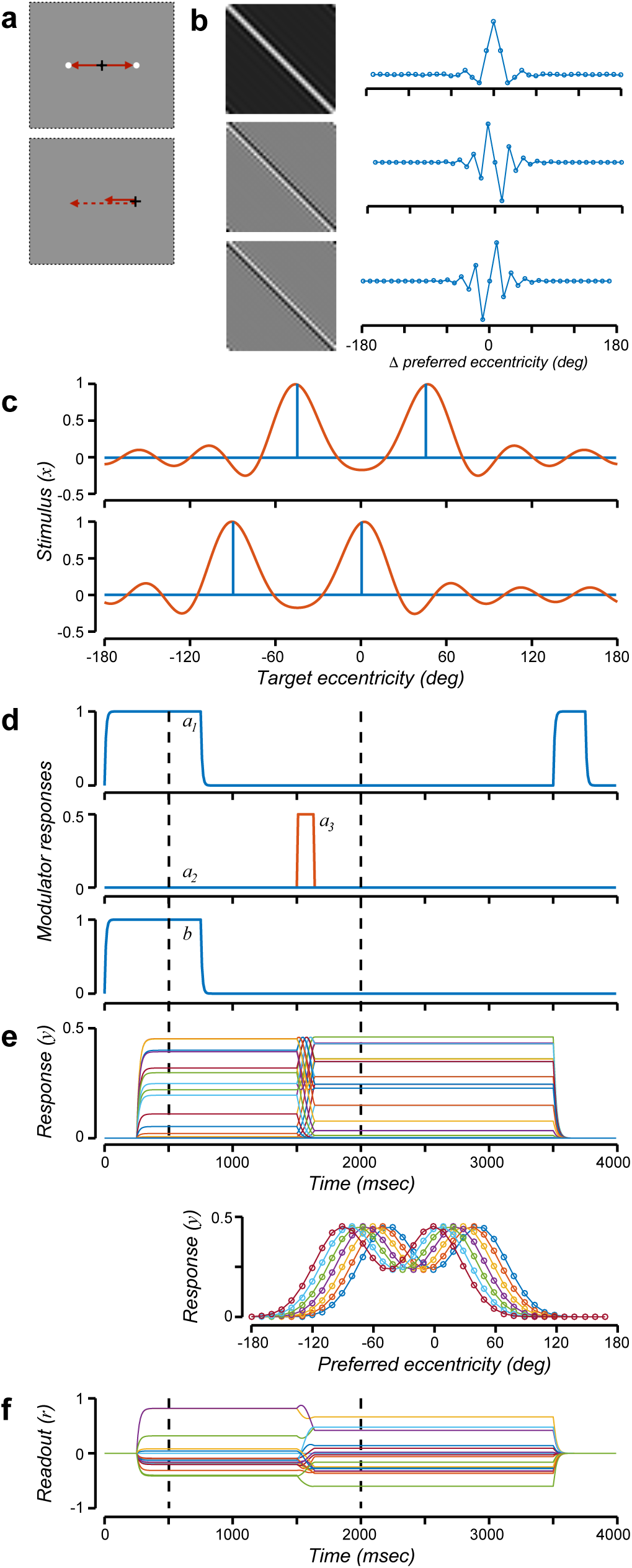
Spatial updating. **a.** Double-step saccade task. Top, targets presented. Bottom, after eye movement to target 1. White dots, targets. Black crosshairs, eye position. Solid red lines, planned eye movements without updating. Dashed red line, planned eye movement after updating. **b**. Recurrent weight matrices. Top row, recurrent weight matrix corresponding to modulator *a*_1_ for maintaining a representation of the target locations. Middle and bottom rows, recurrent weight matrix corresponding to modulators *a*_2_ and *a*_3_ for updating the representation with leftward and rightward eye movements, respectively. **c.** Input stimulus and reconstructed stimulus. Blue, input stimulus (**x**) corresponding to the two target positions. Orange, reconstructed stimulus, computed as a weighted sum of the readout (panel f). Top row, before eye movement to target 1. Bottom row, after eye movement to target 1. **d.** Modulator responses. Top row, *a*_1_. Middle row, *a*_2_ (blue) and *a*_3_ (orange). Bottom row, *b*. **e.** Output responses (**y**). Top, time course of activity, with different colors corresponding to different neurons. Bottom, responses for each of several time points (different colors correspond to different time points) while updating the neural representation of the target locations. **f.** Readout (**r**). Dashed vertical lines in panels d-f correspond to the snapshots in panel a.

The example circuit in **Fig. 6** received two types of inputs: 1) the target locations at the beginning of the trial (**Fig. 6c**, top row, blue), and 2) a corollary discharge of the impending eye movement. The targets were assumed to be along the horizontal meridian of the visual field. There were again 36 neurons, but unlike the preceding examples, each neuron responded selectively to a different eccentricity along the horizontal meridian of the visual field (i.e., degrees of visual angle away from fixation), not different polar angles around fixation at a fixed eccentricity. The encoding matrix **W**_*zx*_ was analogous to that in the preceding examples, but the neurons were selective for target eccentricity instead of polar angle. Readout and reconstruction was the same as that for the sustained activity circuit (**Fig. 2**).

What distinguishes this example circuit from the preceding examples is that there were three recurrent weight matrices (**Fig. 6b**), the first for maintaining a representation of the target locations (**Fig. 6b**, top row), the second for changing the representation with leftward eye movements (**Fig. 6b**, middle row), and the third for changing the representation with rightward eye movements (**Fig. 6b**, bottom row). As in the preceding examples, the modulators were the same for each neuron in the circuit. Consequently, we can modify **Eq. 1**:

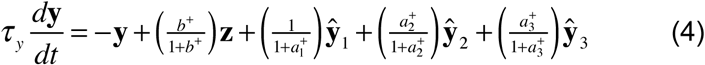

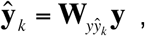

where the subscript *k* indexes over the 3 recurrent weight matrices. The first recurrent weight matrix was identical to that in the sustained activity circuit (**Fig. 2b**). The second recurrent weight matrix was a discrete approximation to the derivative of the responses (see Supplementary Material), and third was the negative derivative matrix (i.e., the second and third recurrent matrices differed from one another by a factor of −1).

The modulators were used to encode and update a representation of the target locations (**Fig. 6d**). As in the preceding examples, the responses followed the input drive at the beginning of the simulated trial because *b* and *a*_1_ were large (=1, corresponding to a short effective time constant). The values of *a*_1_ and *b* then switched to be small (=0, corresponding to a long effective time constant) before the targets were extinguished, so the output responses exhibited sustained activity that represented the original target locations (**Fig. 6c**, top row, orange). The modulator *a*_3_ was non-zero for a period of time beginning just prior to the eye movement (**Fig. 6d**, middle row, orange). The amplitude of *a*_3_ and duration of time during which it was non-zero determined the magnitude of updating, i.e., corresponding to the amplitude of the impending saccade. Finally, the value of *a*_1_ was large (=1, corresponding to a small recurrent gain) at the end of the trial, causing the output responses to be extinguished.

The output responses exhibited a traveling wave of activity during the period of time when the neural representation of the targets was updated (**Fig. 6e**), i.e., another example of sequential activity. The readout (**Fig. 6f**) encoded the two target locations, both before and after updating. The readout, at any time point, was multiplied by a fixed decoding (or reconstruction) matrix to approximately reconstruct the input stimulus (see Supplementary Material). Preceding the eye movement, the original target locations were reconstructed from the readout (**Fig. 6c**, top row, orange curve). After the eye movement, the updated target locations were reconstructed (**Fig. 6c**, bottom row), using the same decoding matrix.

### Manipulation: time warping and time reversal

A challenge for models of motor control is to generate movements at different speeds, e.g., playing the same piece of piano music at different tempos. Likewise, a challenge for models of sensory processing is that perception must be invariant with respect to compression or dilation of temporal signals, e.g., fast vs. slow speech (70). Scaling the time constants of the neurons, all by the same factor, scales the oscillation frequencies by the inverse of that scale factor (**Eq. 3**). This offers a possible mechanism for time warping (71-74). A fixed value for the scale factor would handle linear time-rescaling in which the entire input (and/or output) signal is **b** compressed or dilated by the inverse of the scale factor. A neural circuit might compute a time-varying value for the scale factor, based on the inputs and/or outputs, to handle time-varying time-warping.Here, we offer a different mechanism for time warping (also time reversal), making use of the modulator responses.

An example of open-loop motor control was implemented that enabled time warping and time reversal (**Fig. 7**). The encoding matrix and the recurrent matrix were identical to those in the spatial updating example (**Fig. 6**). The *a*_1_ and *b* modulators were also the same as in the spatial updating example, but the other two modulators *a*_2_ and *a*_3_ were different (**Fig. 7a**). The readout was the same as that in the motor control circuit (**Fig. 5**), summing across the components *r***^Σ^**. The input was chosen to drive all of the eigenvectors with randomly chosen amplitudes and phases. Different values of the *a*_2_ and *a*_3_ modulators generated control signals that were time warped and/or time reversed. Increasing the modulator response from 1 to 5/3 caused the readout to increase in tempo by 25% (compare **Figs. 7b** and **7c**); tempo was proportional to *a*_2_ / (1 + *a*_2_). When *a*_2_ was zero and *a*_3_ was non-zero, then the readout was time reversed (compare **Figs. 7b** and **7d**). A time-varying modulator generated time-varying time-warping (not shown). The circuit exhibited these phenomena because the responses exhibited oscillating traveling waves (see **Fig. 5b**). The readout was a sum of these traveling waves, and the speed of the traveling waves was controlled by the modulators (see Supplementary Materials).

**Figure 7.**
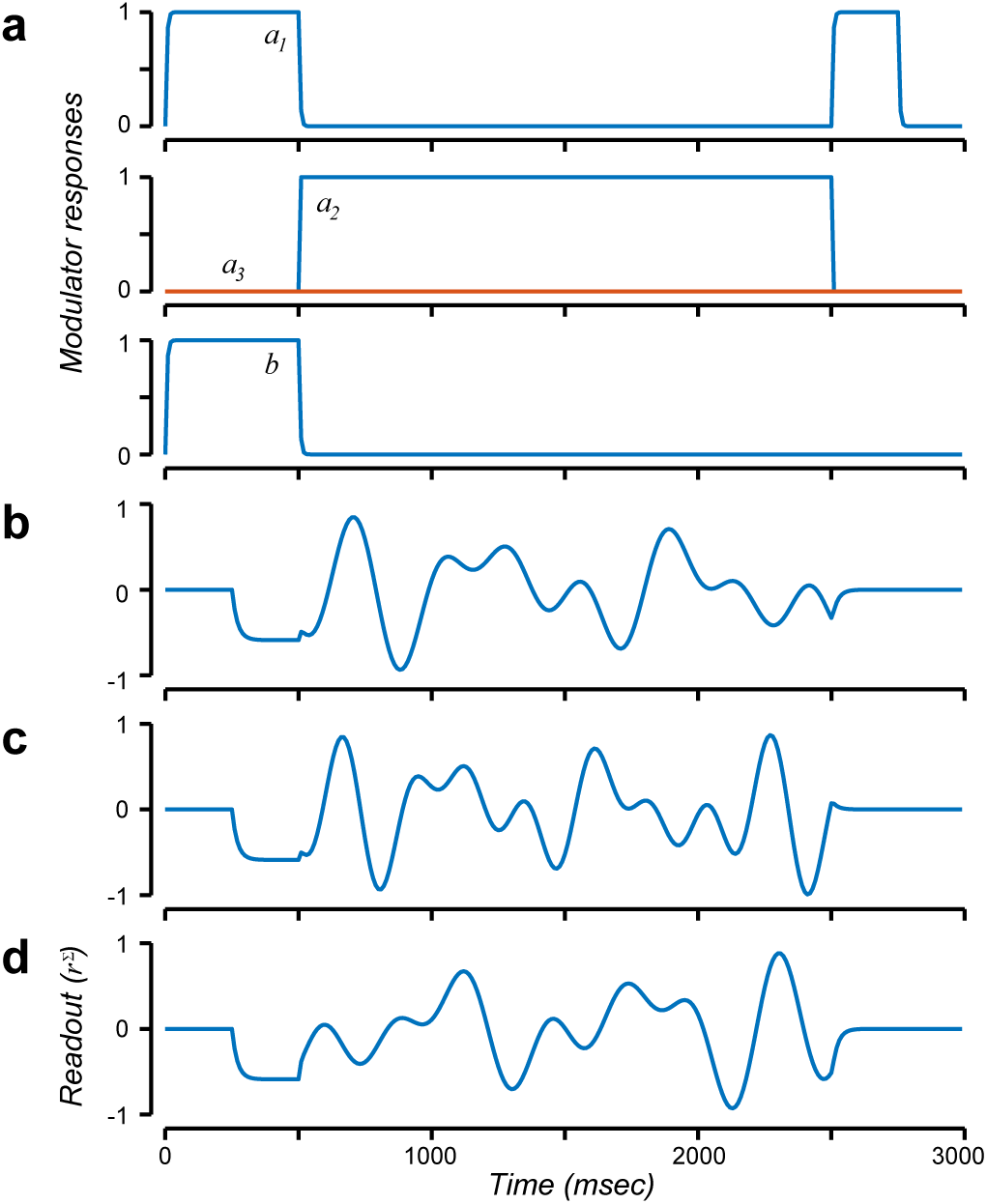
Time warping and time reversal. **a**. Modula**a** tor responses. **b.** Readout for *a*_2_ = 1 and *a*_3_ = 0. **c.** Time-warped readout for *a*_2_ = 5/3 and *a*_3_ = 0. **d.** Time-reversed readout for *a*_2_ = 0 and *a*_3_ = 1.

### Robustness via normalization

The example circuits discussed thus far depended on precisely tuned synaptic weights. The recurrent weight matrices were scaled so that the eigenvalues had real parts no greater than 1. If the recurrent weight matrix has eigenvalues with real parts greater than 1, then the responses are unstable, growing without bound during a delay period. This is a well-known problem for recurrent neural networks (75, 76).

A solution to this problem is to combine ORGaNICs with normalization. The normalization model was initially developed to explain stimulus-evoked responses of neurons in primary visual cortex (V1) (58, 77, 78), but has since been applied to explain neural activity in a wide variety of neural systems (79). The model’s defining characteristic is that the response of each neuron is divided by a factor that includes a weighted sum of activity of a pool of neurons. The model predicts and explains many well-documented physiological phenomena, as well as their behavioral and perceptual analogs.

A variant of ORGaNICS was implemented that used the recurrent modulator (*a*) to provide normalization. The recurrent modulator *a* determined the amount of recurrent gain; it was a particular nonlinear function of the responses *y* instead of the linear function expressed by **Eq. 2** (see Supplementary Materials for details). For an input drive **z** that was constant for a period of time, the output responses achieved a stable state in which they were normalized (see Supplementary Materials for derivation):

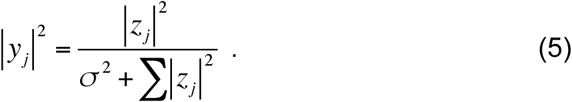

The responses were proportional to the input drive when the amplitude of the input drive was small (i.e., when the sum of the squared input drives was ≪ *σ*). The responses saturated (i.e., leveled off) when the amplitude of the input drive was large (≫ *σ*). The value of *σ* (the semi-saturation constant) determined the input drive amplitude that achieved half the maximum response. In spite of saturation, the relative responses were maintained (see Supplementary Materials for derivation):

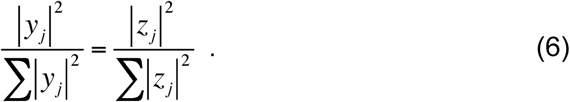

That is, the normalized responses represented a ratio between the input drive to an individual neuron and the amplitude of the input drive summed across all of the neurons. Consequently, the responses of all of the neurons saturated together (at the same input drive amplitude) even though some neurons responded strongly to the target location whereas others did not.

Recurrent normalization made the circuit robust with respect to imperfections in the recurrent weight matrix (**Fig. 8**). Without normalization, responses depended critically on fine tuning. For example, we used the sustained activity circuit (**Figs. 1-2**), but scaled the recurrent weight matrix by a factor of 1.02. The responses were unstable, growing without bound (**Fig. 8a**). Including normalization automatically stabilized the activity of the circuit (**Fig. 8b**). The increases in activity evoked by the recurrent weight matrix (with the largest eigenvalues = 1.02) were countered by normalization such that the total activity in the circuit was roughly constant over time (||**y**||^2^ ≈ 1). The ratios of the responses were maintained (**Eq. 6**), enabling an accurate readout, throughout the delay period. Analogous results (not shown) were obtained with the other example circuits described above, including those that exhibited oscillatory and sequential dynamics, because the normalization depends on the squared norm of the responses which was constant over time during the delay period for each of these example circuits.

**Figure 8.**
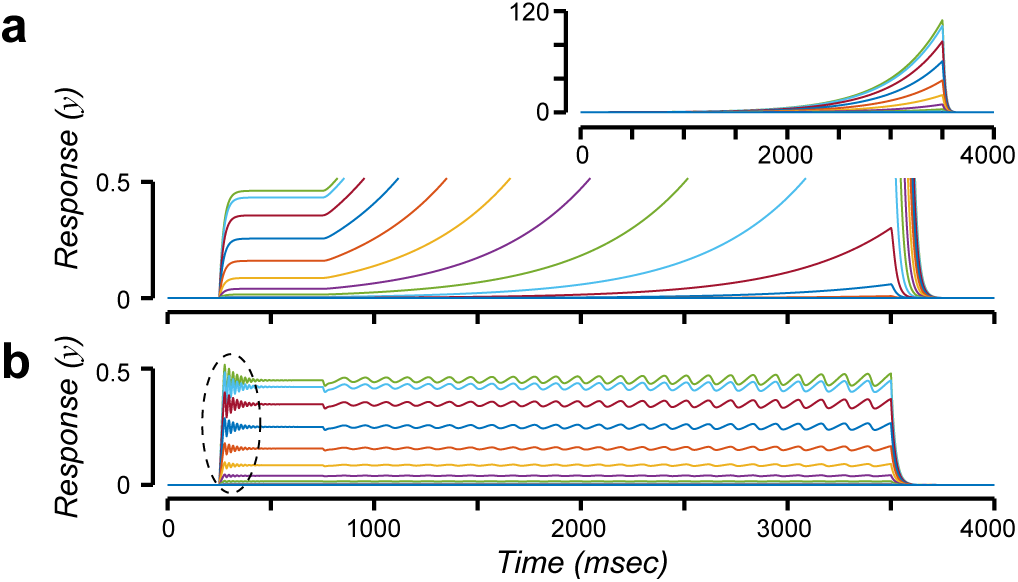
Robustness and normalization. **a.** Output responses (**y**), corresponding to the sustained activity circuit depicted in Figs. 1-2, but with the recurrent weight matrix scaled by a factor of 1.02. Each color corresponds to a different neuron. Inset, full range of responses on expanded (240x) ordinate. **b.** Output responses with normalization. Dashed oval, high frequency, coherent, synchronized oscillations following target onset.

The normalized responses exhibited high frequency, coherent, synchronized oscillations following target onset (**Fig. 8b**, dashed oval). There are two nested recurrent circuits in ORGaNICs: 1) the recurrent drive, and 2) the multiplicative modulators. The high-frequency oscillations emerged because of the inherent delay in the second of these recurrent circuits, i.e., because of the multiplicative modulator underlying normalization. The oscillation frequency depended on the membrane time constants. For the time constants used for **Fig. 8**, the responses exhibited oscillations in the gamma frequency range. Different intrinsic time constants yielded different oscillation frequencies. The oscillation frequency would have depended also on axon length if we were to include conduction delays. The responses exhibited lower frequency oscillations during the delay period (**Fig. 8b**). These lower frequency oscillations emerged because of the recurrent drive in combination with normalization; the recurrent weight matrix was scaled to have eigenvalues greater than 1 which drove the responses to increase over time but this increase was countered by normalization.

### Biophysical implementation

A possible biophysical implementation of OR-GaNICs is depicted in **Fig. 9**. The key idea is that the two terms corresponding to the input drive and recurrent drive are computed in separate dendritic compartments of a cortical pyramidal cell. The model comprises 3 compartments for the soma, the apical dendrite, and the basal dendrite. Each compartment is an RC circuit with a variable-conductance resistor and a variable current source. The capacitors represent the electrical capacitance of the neural membrane. The two fixed-conductance resistors (*R*_*a*_ and *R*_*b*_) represent the resistances between the compartments (i.e., along the dendritic shafts). The membrane potentials *v*_*s*_, *v*_*a*_, and *v*_*b*_ correspond to the soma, the apical dendrite, and the basal dendrite, respectively. Some of the synaptic inputs are modeled as conductance changes (*g*_*va*_, *g*_*vb*_, *g*_*vs*_) while others are approximated as currents (*I*_*s*_, *I*_*a*_, *I*_*b*_). To implement ORGaNICs with this pyramidal-cell model, we specified the synaptic inputs (*I*_*s*_, *I*_*a*_, *I*_*b*_, *g*_*va*_, *g*_*vb*_, and *g*_*vs*_) to each neuron in terms of its input drive (*y*), recurrent drive (*ŷ*), and modulators (*a* and *b*). We also presumed that the output firing rate of a neuron was well-approximated by halfwave rectification of the membrane potential, and that negative values (corresponding to hyperpolarlization of the membrane potential *v*_*s*_) were represented by a separate neuron that received the complementary synaptic inputs (identical for *g*_*va*_ and *g*_*vb*_, and opposite in sign for *I*_*s*_, *I*_*a*_, and *I*_*b*_), analogous to ON-and OFF-center retinal ganglion cells. Then the steady-state value for the somatic membrane-potential (i.e., when the synaptic inputs are constant) is:

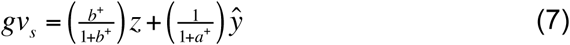

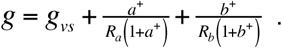

where *g* is the total synaptic conductance (for derivation and detailed description of implementation, see 56).

**Figure 9.**
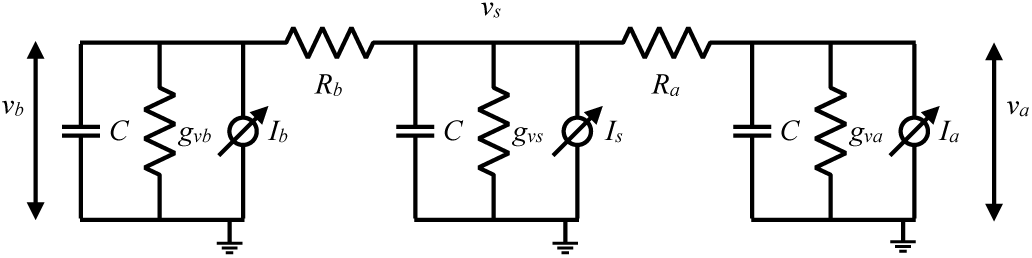
Biophysical implementation. Electrical-circuit model of a pyramidal cell with separate RC circuit compartments for the soma, the apical dendritic tree, and the basal dendritic tree. *g*_*va*_, *g*_*vb*_, shunting synapses represented by variable conductance resistors. *I*_*a*_, *I*_*b*_, synaptic input represented by variable current sources. *v*_*s*_, somatic membrane potential. *v*_*a*_, *v*_*b*_, membrane potentials, respectively, in apical and basal dendrites. *R*_*a*_, *R*_*b*_, fixed-conductance resistors between compartments (i.e., along the dendritic shafts). *C*, membrane capacitance.

The steady-state membrane potential (**Eq. 7**) is a weighted sum of the input drive and recurrent drive, modulated by *a* and *b*, respectively, and then scaled by the total somatic conductance. This is identical to the steady-state response of ORGaNICs (compare **Eq. 7** with **Eq. 1**) when the total somatic conductance is *g* = 1. There are a variety of combinations of the various parameters of this biophysical model for which the total somatic conductance is approximately equal to 1 (56). Two particular interesting special cases correspond to when the modulators are both on (i.e., equal to 1, such the responses are dominated by the input drive), and when the modulators are both off (i.e., equal to 0, during a delay period).

This is, of course, a simplified model of pyramidal cells. First, it has been argued that shunting inhibition does not yield division (80); *in vivo* neurons are rarely at their resting potential because of spontaneous background activity so a shunting synapse (assumed to be a chloride channel) predominantly causes hyperpolarization rather than division. However, exact division can be implemented with two synaptic conductances, one excitatory and one inhibitory, that increase (and decrease) in proportion (78). And there is experimental evidence that cortical circuits are capable of divisive suppression (81, 82). Second, there is no leak conductance in the dendrites. We can think of *g*_*vs*_ = 1 as the somatic leak conductance, but the model (as expressed above) has no dendritic leak conductances. If we were to add dendritic leak conductances, then the responses would decay during a delay period even when the modulators = 0. One could compensate for this decay by scaling the recurrent weight matrix to have eigenvalues larger than one. Third, the input drive and recurrent drive in the model are mediated by synaptic currents, not conductance changes. A push-pull arrangement of synaptic inputs can act like a current source (78). Doing so necessitates a high level of spontaneous activity so that increases in excitation are met with equal decreases in inhibition, and vice versa. But spontaneous activity in most cortical areas, although non-zero, is generally low. Instead, synaptic inputs could approximate current sources when the membrane potential remains far from the (excitatory and inhibitory) synaptic reversal potentials.

## Discussion

Sustained delay-period activity and sequential activity are opposite sides of the same coin. ORGaNICs, a straightforward extension of leaky neural integrators and neural oscillators, provide a unified theoretical framework for sustained activity (**Fig. 2**), oscillatory activity (**Fig. 3**), and sequential activity (**Fig. 4**), just by changing the recurrent weight matrix. The responses can be read out at any time during the delay period in spite of the complex dynamics. Indeed, we assert that complicated dynamics is the norm, to support manipulation as well as maintenance (e.g., **Fig. 6**).

In addition, ORGaNICs can be used to generate motor control signals, with the very same circuits used to model working memory, just by changing the readout. The circuits convert spatial patterns of input (premotor) activity to temporal profiles of output (motor control) activity. Different spatial patterns of premotor activity evoke motor control outputs with different temporal response dynamics (e.g., as in **Figs. 5** and **7**), and the modulators provide a means for manipulating (time warping and time reversal) the dynamics (**Fig. 7**).

ORGaNICs are applicable also to models of sensory integration (e.g., integrating corollary discharge in **Fig. 6**) and sensory processing (e.g., with normalization as in **Fig. 8**). ORGaNICs can be stacked in layers such that the inputs to one ORGaNIC are the outputs from one or more other ORGaNICs. Particular stacked architectures encompass convolutional neural nets (i.e., deep nets) as a special case: specifically when the encoding/embedding weight matrices are convolutional and when the modulators are large (*a*_*j*_ = *b*_*j*_ ≫ 0) such that the output responses from each layer are dominated by the input drive to that layer. Consequently, working memory, motor control, sensory processing, (including prediction over time, see Supplementary Materials and Ref. 56), and possibly other cognitive functions (in addition to working memory, such as cognitive control, e.g., controlling attention) may all share a common canonical computational foundation.

There is considerable flexibility in the formulation of ORGaNICs, with different variants corresponding to different hypothesized neural circuits with different functions (see Supplementary Materials for examples). ORGaNICs also offer computational advantages compared to varieties of LSTMs that are commonly used in AI applications (see Supplementary Materials and Ref. 56).

The literature on recurrent neural networks includes, of course, various complementary approaches that have each achieved some of the same goals (83-95). But none of the previously published models offer the same breadth of application of ORGaNICs. With ORGaNICs, we show that a single unified circuit architecture captures key phenomena of sensory, motor, and cognitive function. Unlike linear recurrent neural networks, the modulators in ORGaNICs introduce a nonlinearity (analogous to the gates in LSTMs) that can perform multiple functions including handling long-term dependencies and providing robustness via normalization (see below). Nor does the previously published literature provide the same depth of understanding. Unlike most nonlinear recurrent neural nets, ORGaNICs, are mathematically tractable during periods of time when the modulators are constant, i.e., during each successive phase of a behavioral task (see Supplementary Materials for derivations). We can derive concrete, quantitative predictions that can be fit to experimental measurements. For example, the responses of the normalization circuit follow the normalization equation (**Eq. 5**) exactly. Consequently, this circuit makes predictions that are identical to those of the normalization model, thereby preserving all the desirable features of that model, which has been fit to hundreds of experimental data sets. Unlike black-box machine learning approaches, ORGaNICs provide insight; for example, we understand exactly when and how it is possible to reconstruct an input by reading out the responses during the delay period of a working memory task, and how to generate motor control signals with complex dynamics (see Supplementary Materials for derivations). Machine learning algorithms are particularly useful for computing solutions to optimization problems (e.g., model fitting via gradient descent), and we plan to use a machine learning implementation of ORGaNICs to fit experimental data. Machine learning approaches can also provide inspiration for neuroscience theories (and vice versa), like the links presented here between ORGaNICs and LSTMs. Left open in the current paper is how to learn the weights in the various weight matrices in ORGaNICs (see Supplementary Materials). We engineered the weights to demonstrate the computational capabilities of this theoretical framework, and to illustrate that the theory can reproduce neurobiological phenomena. Some of the previously published literature (cited above) focuses on learning. But having the right circuit architecture is a prerequisite for developing of an accurate model of learning.

We posit that the most important contribution of ORGaNICs is the conceptual framework, a canonical computational motif based on recurrent amplification, gated integration, reset, and controlling the effective time constant. Rethinking cortical computation in these terms should have widespread implications.

Why do some neural circuits exhibit sustained activity while others exhibit sequential activity, and what are the relative advantages or disadvantages of each? Sustained activity circuits are useful for short-term memory (i.e., maintenance), but not for other cognitive functions (i.e., that require manipulation and control). For sustained activity circuits, a simple linear readout of the responses can be used to reconstruct the input drive (and to approximately reconstruct the input stimulus), at any point in time during a delay period (**Fig. 2**). In addition, sustained activity circuits are likely to be more robust than sequential activity circuits, because all of the components share the same dynamics. Sequential activity circuits, on the other hand, offer much more flexibility. The same circuit, with the same fixed recurrent weight matrix and the same fixed encoding matrix, can support multiple different functions just by changing the readout. For example, the sequential activity circuit (**Fig. 4**) and the motor control circuit (**Fig. 5**) were identical except for the readout. For the sequential activity circuit (**Fig. 4**), a (nonlinear) modulus readout generated an output that was constant over time (i.e., to support maintenance). For the motor control circuit (**Fig. 5**), a linear readout was used to generate control signals as sums of (co-)sinusoidal basis functions with various different frequencies and phases. Likewise, the spatial updating circuit (**Fig. 6**) and the time warping / time reversal circuit (**Fig. 7**) were identical. This circuit can be used to perform working memory (maintenance and manipulation), and the same circuit (without changing the encoding or recurrent weights) can be used to execute movements with complex dynamics. One way to implement this, for example, would be to have two different brain areas with stereotypical intrinsic circuitry (i.e., identical recurrent weights) that support two different functions with different readouts. Indeed, there is experimental evidence that different brain areas support different functions with similar circuits, e.g., parietal areas underlying working memory maintenance and PFC areas underlying motor planning (96). Alternatively, the output from a single circuit could go to two different brain areas, one of which performs the 1^st^ readout and the other of which performs the 2^nd^ readout. Or a single brain area might switch between two different readouts (e.g., using a gating mechanism analogous to that in ORGaNICs), corresponding to different behavioral states, without changing the intrinsic connectivity within the circuit. This makes biological sense. Rather than having to change everything (the encoding matrix, the recurrent matrix, the modulators, and the readout), you need only change one thing (the readout matrix) to enable a wide variety of functions. This is not possible with recurrent weight matrices that exhibit sustained activity, simply because there is only a single mode of dynamics (constant over time).

ORGaNICs offer a theoretical link between sequential activity and internal models. ORGaNICs can be used to maintain and manipulate information over time. For example, one of our example circuits encoded a representation of a pair of targets and then updated the representation (of the remembered locations) coincident with an eye movement (**Fig. 6**). This process of updating the representation of a visual target has been called remapping (97-99). The responses during the update exhibited a traveling wave of activity (i.e., sequential activity), commensurate with measurements of neural activity during remapping (100). We can think of this as an internal model (also called a forward model) of the movement. Internal models are hypothesized to predict the result of a movement plan and to explain how an organism discounts sensory input caused by its own behavior (101-105). Our simulation results are also reminiscent of an ostensibly different class of phenomena: internally-generated sequential activity in parietal cortex, prefrontal cortex, and hippocampus during navigation, motor planning, and episodic recall (21-23, 106). We hypothesize that these forms of sequential activity can be explained by the same gated integration computation that we used for spatial updating, i.e., updating an internal model.

Motor control may share a common computational foundation with working memory (and possibly other cognitive functions), performed with similar circuits. Open-loop (i.e., ballistic) actions can be performed by initializing the state of an ORGaNIC (i.e., with the modulators turned on) with a spatial pattern of premotor activity, and then switching the modulators to let the dynamics play out, thereby converting the spatial pattern of premotor activity to a temporal profile of motor control activity. This is commensurate with experimental evidence demonstrating that motor preparation sets the initial conditions for subsequent neural dynamics underlying movement execution (66). Closed-loop actions may also be performed with the same computation and circuit, in which the input drive reflects sensory feedback during the movement.

The normalization circuit (**Fig. 8**) is consonant with the idea that normalization operates via recurrent amplification, i.e., that weak inputs are strongly amplified but that strong inputs are only weakly amplified. Several hypotheses for the recurrent circuits underlying normalization have been proposed (78, 79, 107-112), but most of them are inconsistent with experimental observations suggesting that normalization is implemented via recurrent amplification (113-118). ORGaNICs offer a family of dynamical systems models of normalization, each of which comprises coupled neural integrators to implement normalization exactly; when the input drive is constant over time, the circuit achieves an asymptotic stable state in which the output responses follow the normalization equation (**Eq. 5**).

ORGaNICs provide a theoretical framework for dimensionality-reduction data-analysis methods. Sensory, cognitive, and motor functions depend on the interactions of many neurons. It has become increasingly popular to use dimensionality-reduction algorithms to analyze measurements of neural activity recorded simultaneously in large numbers of neurons (119, 120). These data-analysis algorithms presume that the neural activity over time is confined to a low-dimensional manifold, much lower than the number of neurons recorded. The simplest of these algorithms use principal components analysis or factor analysis, thereby presuming that the neural activity is confined to a linear subspace. The simulated responses of the ORGaNICs are indeed confined to a linear subspace at any moment in time, but the subspace changes over time with changes in the state. The responses are in the subspace of the encoding matrix when the modulators are both large. The responses are in the subspace of the eigenvectors of the recurrent weight matrix (with corresponding eigenvalues that have real parts = 1) when modulators are both 0 for long enough such that the other components decay. The responses can be in any of an infinite number of subspaces for intermediate values of the modulators and with non-zero inputs.

Our theory offers a generalization of the notion of E:I balance. The stability of ORGaNICs (and related neural integrator circuits) depends on a combination of the recurrent weight matrix and the relative values of the intrinsic time constants. For example, a differential delay between excitatory and inhibitory neurons can compensate for an imbalance in the excitatory versus inhibitory synaptic weights, and vice versa (56).

The theory predicts that changes in state are associated with measurable changes in response dynamics. Of particular interest is the change in intrinsic oscillation frequency evident in the normalization circuit (**Fig. 8B**). That circuit exhibited high frequency, coherent, synchronized oscillations following target onset, and lower frequency oscillations during the delay period. The change in oscillation frequency corresponded to a change in state, induced by changing the modulator responses. This is commensurate with experimental observations that different brain states are associated with particular oscillation frequencies (32, 33, 121-127). The high-frequency oscillations emerged because of the multiplicative modulator underlying normalization; the oscillation frequency depended on the membrane time constants, and would have depended also on axon length if we were to include conduction delays. The emergence of high-frequency oscillations in our normalization circuit dovetails with observations that gamma oscillations are linked to normalization (128-134). But this multiplicative (i.e., nonlinear) process is different from most previous models of gamma oscillations (110, 131, 135-139), which fundamentally rely on linear recurrent neural oscillators (see Supplementary Materials for a primer on neural oscillators).

The modulators perform multiple functions and can be implemented with a variety of circuit, cellular, and synaptic mechanisms. The time-varying values of the modulators, **a** and **b**, determine the state of the circuit by controlling the recurrent gain and effective time-constant of each neuron in the circuit. The modulators, and consequently the dynamically-changing state, depend on the current inputs and the current outputs, which in turn depend on past inputs and outputs. The multiple functions of the modulators include normalization (**Fig. 8**), sustained activity (**Fig. 2**), controlling pattern generators (**Fig. 5**, **Fig. 7**, and **Fig. S2**), gated integration/updating (**Fig. 6**), time warping and time reversal (**Fig. 7**), reset (**Figs. 2-8**), and controlling the effective time constant (**Fig. S1**). Each neuron may have multiple modulators to perform combinations of these functions (e.g., **Fig. 6**). Some of the modulator functions need to be fast and selective (e.g., normalization), likely implemented in local circuits. A variety of mechanisms have been hypothesized for adjusting the gain of local circuits (78, 140, 141). Some modulator functions might depend on thalamocortical loops (24, 142-144). Other modulator functions are relatively non-selective and evolve slowly over time, and may be implemented with neuromodulators (145-147).

There is a critical need for developing behavioral tasks that animal models are capable of learning, and that involve both maintaining and manipulating information over time. ORGaNICs (and LSTMs) manage long-term dependencies between sensory inputs at different times, using a combination of gated integration (e.g., when *b*_*j*_ > 0 and *a*_*j*_ = 0) and reset (e.g., when *a*_*j*_ > 0 and *b*_*j*_ = 0). Typical delayed-response tasks like the memory-guided saccade task are appropriate for studying what psychologists call “short-term memory”, but they are weak probes for studying working memory (148-151), because those tasks do not involve manipulation of information over time. Behavioral tasks that are popular in studies of decision-making involve integration of noisy sensory information (152-155) or integration of probabilistic cues (156). Variants of these tasks (36, 157) might be used to test the gated integration and reset functionality of ORGaNICs. The anti-saccade task (158-162) and the double-step saccade task (163-167) might also be used, with delay periods, to test the theory and to characterize how cortical circuits manage long-term dependencies.

Finally, the theory motivates a variety of experiments, some examples of which are as follows. 1) The theory predicts that the modulators change the effective time constant and recurrent gain of a PFC (or PPC) neuron. Experimental evidence suggests that the modulatory responses corresponding to the modulators, **a** and **b**, are computed in the thalamus (2, 24, 142). Consequently, manipulating the responses of these thalamic neurons (e.g., via optogenetics) should have a particular impact on both the time constant and recurrent gain of cortical neurons. 2) The specific biophysical implementation (**Fig. 9**) predicts that the soma and basal dendrites share input drive, but with opposite sign. This would, of course, have to be implemented with inhibitory interneurons. 3) The theory predicts that a dynamic time-varying pattern of neural activity (e.g., sequential activity) can nonetheless encode information that is constant over time. In such cases, it ought to be possible to read out the encoded information using a fixed decoder in spite of dynamic changes in neural activity. 4) As noted above, variants of sensory integration tasks might be used to test the gated integration and reset functionality of ORGaNICs, and variants of the anti-saccade and double-step saccade tasks might also be used, with delay periods, to characterize how cortical circuits manage long-term dependencies.

## Acknowledgements

Funding: None

Special thanks to Mike Landy, Mike Halassa, Eero Simoncelli, and Charlie Burlingham for comments and discussion.

## Supplementary Material

### Primer on leaky neural integrators and neural oscillators

A leaky neural integrator (**Fig. S1**) corresponds to a special case of ORGaNICs. For this special case, the modulators *a* = *b* are equal to one another and constant over time, and the recurrent weight matrix equals the identity matrix. A simple example with a single neuron is expressed as follows:

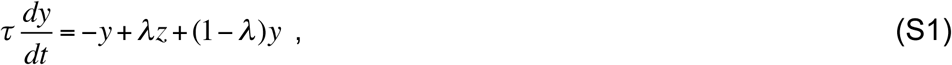

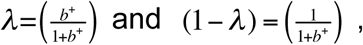

i.e.,

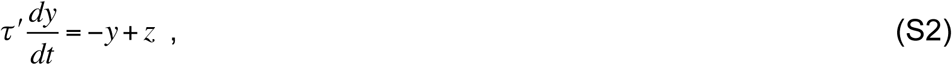

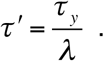

where *τ* is the intrinsic time-constant and *τ′* is the effective time-constant. For this simple special case, the neuron acts like a shift-invariant linear system, i.e., a recursive linear filter with an exponential impulse response function. If the input drive *z* is constant over time, then the responses *y* exhibit an exponential time course with steady state *y* = *z*, and time constant *τ′* (**Fig. S1**). This special case reveals how *λ*, and consequently the modulators, *a* and *b*, determine the effective time-constant of the leaky integrator.

**Figure S1.**
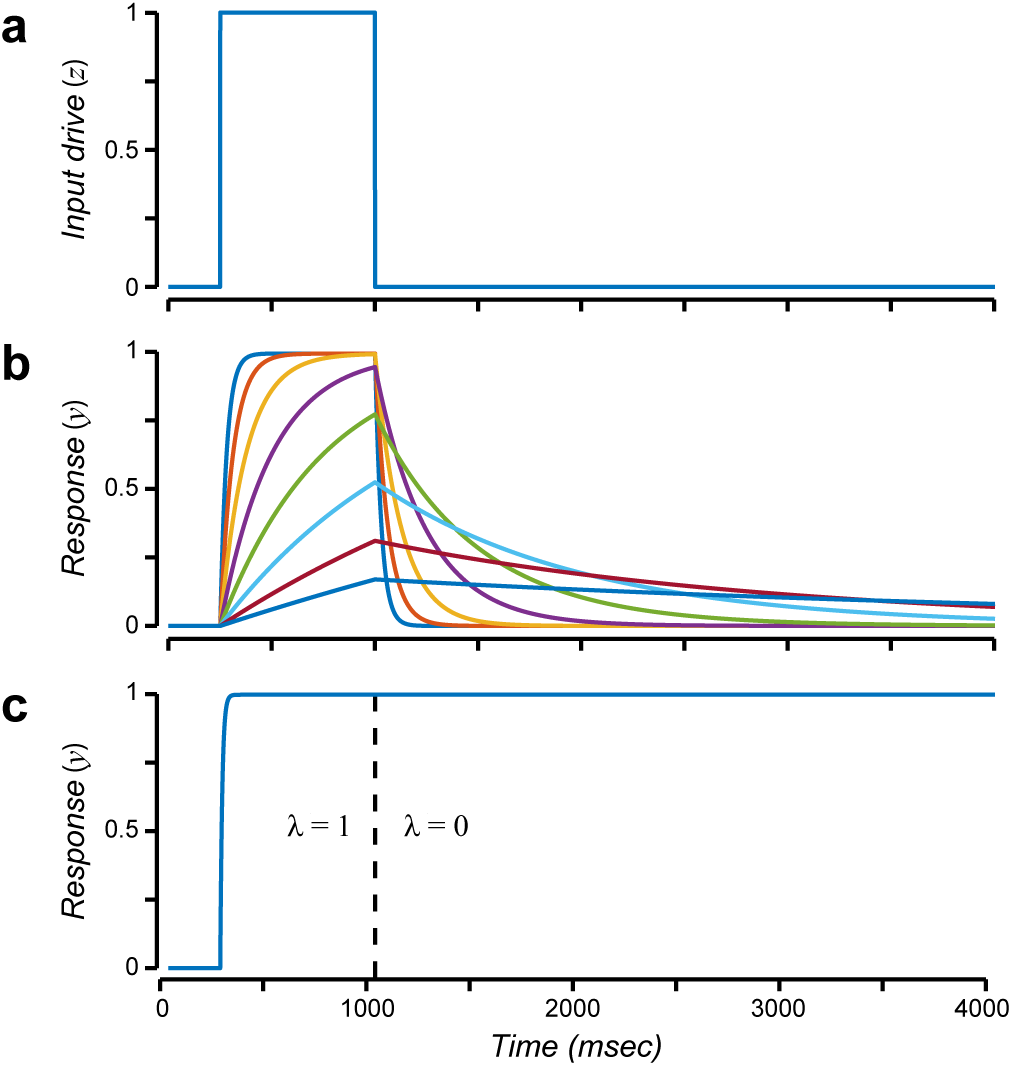
Leaky neural integrator (Eq. S1). **a.** Input drive (*z*) over time. **b.** Output responses (*y*) when the modulator (*λ*) is constant over time. Different colors correspond to different values of *λ*. **c.** Output responses (*y*) corresponding to time-varying modulator (*λ* = 1 for *t* < 1000 and *λ* = 0 for *t* > 1000).

The simple form in **Eq. S1** is problematic because it allows negative firing rates (the value of *y* could be positive or negative depending on the input drive). To fix that, we use a complementary pair of neurons (analogous to ONand OFF-center retinal ganglion cells) that receive complementary copies of the input drive, *z* and -*z*, and for which the firing rates are a halfwave-rectified copy of the underlying membrane potential fluctuations (optionally with a scale factor to convert from mV to spikes/sec):

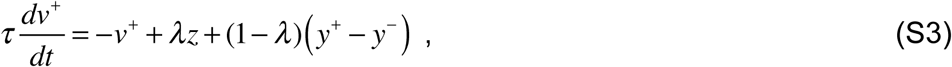

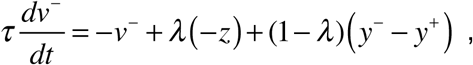

where

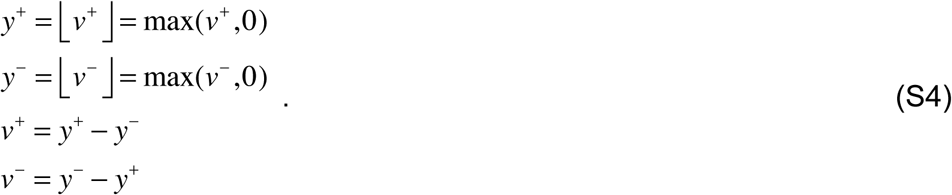

Here, *v*^*+*^ and *v*^*-*^ are the membrane potential fluctuations (and also the recurrent drive) of the two neurons, *z* is the input drive, and *y*^*+*^ and *y*^*-*^ are is the output firing rates.

A leaky neural integrator with multiple neurons comprises a circuit such that the output responses of each neuron depend on a recurrent weighted sum:

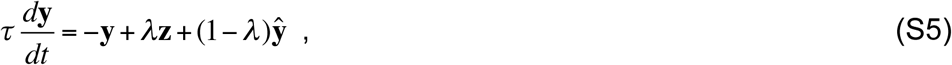

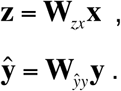

We use boldface lowercase letters to represent vectors and boldface uppercase to denote matrices. The variables (**y**, **ŷ**, **z**, **x**) are each functions of time. The time-varying output responses are represented by a vector **y** = (*y*_*1*_, *y*_*2*_,…, *y*_*j*_,…, *y*_*N*_) where the subscript *j* indexes different neurons in the network. The time-varying input drive is represented by another vector **z** = (*z*_*1*_, *z*_*2*_,…, *z*_*j*_,…, *z*_*N*_). The input stimulus is a continuous function *x*(*θ,t*) where *θ* parameterizes the stimulus space (e.g., polar angle and/or eccentricity in the visual field). The input stimulus is sampled to be represented by another time-varying vector **x** = (*x*_*1*_, *x*_*2*_,…, *x*_*j*_,…, *x*_*M*_), where *x*_*j*_(*t*) = *x*(*θ*_*j*_,*t*) and *θ*_*j*_ are the locations of the samples. The input drive *z*_*j*_ to each neuron is a weight sum of the input **x**, and the weights are given by the encoding matrix weight matrix **W**_*zx*_ (second line of **Eq. S5**). The recurrent drive *ŷ*_*j*_ to each neuron is a weighted sum of the outputs, and the weights are given by the recurrent weight matrix **W**_*ŷy*_ (third line of **Eq. S5**). We can use the same trick as above (**Eqs. S3-S4**) to ensure non-negative firing rates.

A neural oscillator (**Fig. S2**) corresponds to a special case of a leaky neural integrator in which the responses and recurrent weights are complex-valued. Specifically:

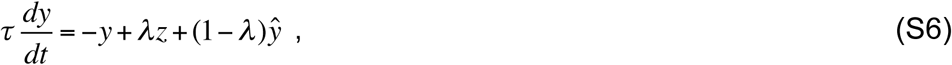

**Figure S2.**
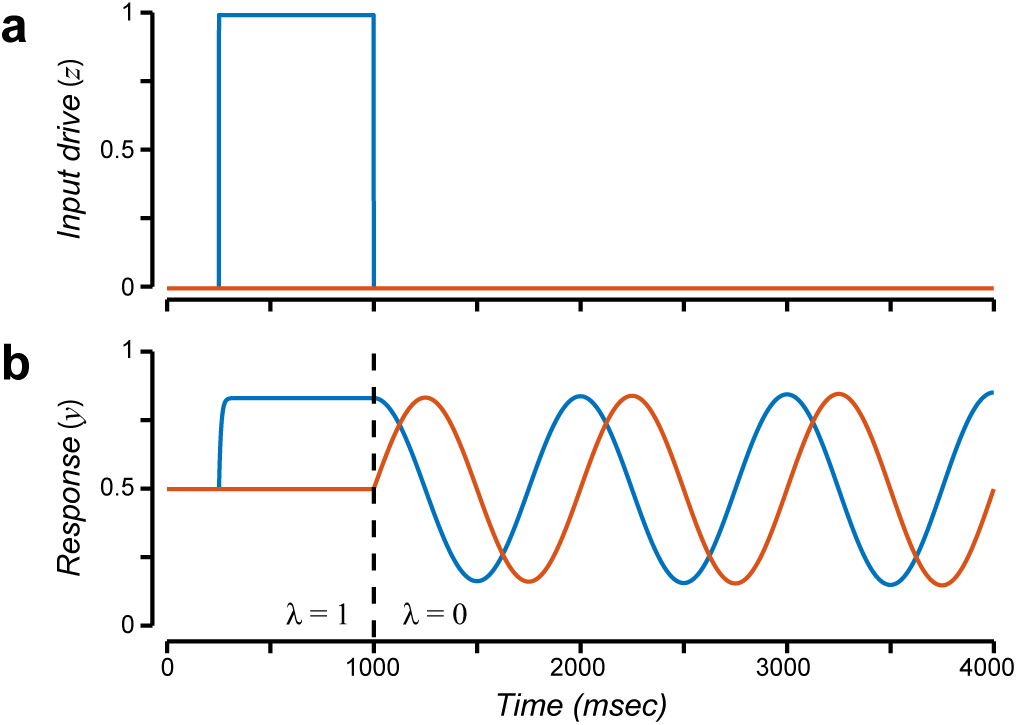
Neural oscillator (Eq. S7). **a.** Input (*z*) over time. Blue, real-part. Orange, imaginary part. Output responses (*y*) corresponding to timevarying modulator (*λ* = 1 for *t* < 1000 and *λ* = 0 for *t* > 1000). Blue, real-part. Orange, imaginary part.

**Figure S3.**
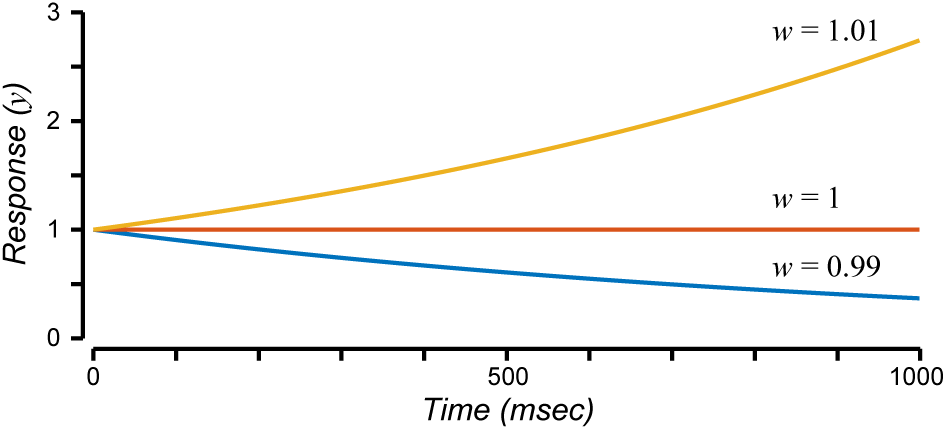
Stability of sustained activity. Responses when *λ* = 0 for 3 different values of the recurrent weight. Yellow, responses grow without bound when recurrent weight is greater than 1. Blue, responses decay to zero when recurrent weight is less than 1. Orange, responses are constant over time when recurrent weight is equal to 1.

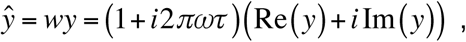

The input drive *z*, responses *y*, and recurrent drive *ŷ* are complex numbers, *w* is the complex-valued recurrent weight, and *ω* is the oscillation frequency. The complex-number notation is just a notational convenience. The complex-valued responses may be represented by a pair of neurons, and the complex-valued weight may be represented by pairs of synaptic weights that are matched to one another in the two neurons:

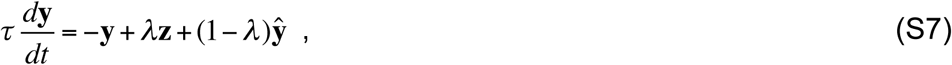

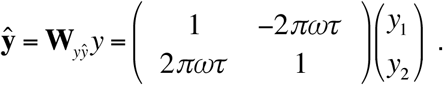

The values of *y*_1_ and *y*_2_ correspond, respectively, to the real and imaginary parts of the complex-valued responses. Intuitively, the responses oscillate because the recurrent weight matrix is a rotation matrix in the limit when the rotation angle is small, i.e., cos(*θ*) = 1 and sin(*θ*) = *θ* when *θ*⟶0. If, at one instant in time, **y** = (1 0)^t^, then an instant later the responses will have changed akin to rotating slightly around the unit circle.

The dynamics of the responses depend on the recurrent weight matrix (**Fig. S3**). This is particularly important when *λ* = 0 (corresponding to the sustained delay-period activity in **Fig. S1** or the period of stable oscillations in **Fig. S2**). If the weights are too small then the responses decay over time. If the weights are too large then the responses grow without bound. A simple is example is given by a circuit with 3 neurons and a diagonal recurrent weight matrix:

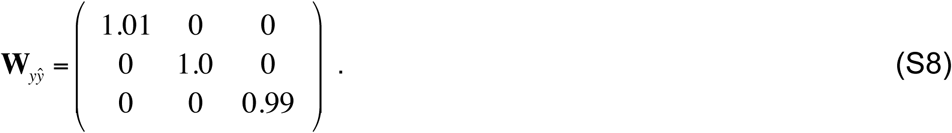

Rewriting **Eq. S5** for the special case of a diagonal weight matrix:

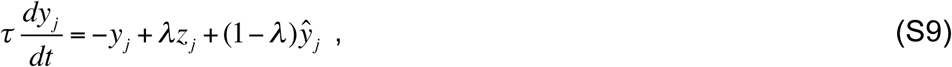

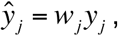

where the recurrent weights *w*_*j*_ are the elements along the diagonal of the recurrent weight matrix **W**_*ŷy*_. When of *λ* = 0, this equation simplifies further:

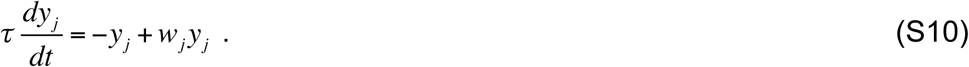

For *w*_*j*_ = 1, the responses are constant over time (the derivative in **Eq. S10** is 0). For *w*_*j*_ > 1, the response grow over time. And for *w*_*j*_ < 1, the response decay over time.

In general, for an arbitrary recurrent weight matrix, the dynamics of the responses depend on the eigenvalues and eigenvectors of the recurrent weight matrix. When the eigenvectors and eigenvalues of the recurrent weight matrix are composed of complex values, the responses exhibit oscillations. For example, the recurrent weight matrix in the neural oscillator example (**Fig. S2**) is an anti-symmetric, 2×2 matrix, with complex-valued eigenvalues and eigenvectors (**Eq. S7**). The real-parts of the eigenvalues determine stability. In this case, the real parts of the eigenvalues are equal to 1 (the weight matrix was in fact scaled so that the eigenvalues have real parts that were equal to 1). The corresponding eigenvectors define a coordinate system (or basis) for the responses. The responses during the period of stable oscillations (when *λ* = 0) are determined entirely by the projection of the initial values (the responses just before the input was turned off) onto the eigenvectors. Eigenvectors with corresponding eigenvalues that have real parts equal to 1 are sustained. Those with eigenvalues that have real parts less than 1 decay to zero (smaller eigenvalues decay more quickly). Those with eigenvalues that have real parts greater than 1 grow without bound (which is why the weight matrix was scaled so that the largest eigenvalues = 1). The imaginary parts of the eigenvalues of the recurrent weight matrix (in this example equal to 2π*ωτ*) determine the oscillation frequencies (*ω* in this example).

### Sustained activity (Fig. 2): encoding, recurrent, readout and reconstruction matrices

The recurrent weight matrix **W**_*ŷy*_ for this circuit was a 36×36 matrix, designed based on quadrature-mirror filter wavelets (1, 2). Quadrature-mirror filters are mutually orthogonal when shifted or scaled by factors of 2. Consequently, the rows of **W**_*ŷy*_ were shifted copies of one another (i.e., **W**_*ŷy*_ was convolutional), but all the even (or odd) rows of **W**_*ŷy*_ were mutually orthogonal so that that the 36 rows of the recurrent weight matrix, corresponding to the 36 neurons in the circuit, spanned a 19-dimensional subspace.

The steady-state responses during the delay period depended on the dot products of the initial responses and the eigenvectors of the recurrent weight matrix **W**_*ŷy*_ with corresponding eigenvalues equal to 1:

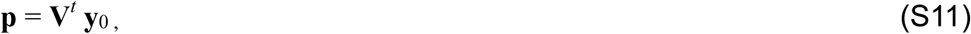

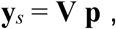

where **y**_*s*_ is the vector of steady-state responses, **y**_0_ is the vector of initial values at the beginning of the delay period, and **V** is a 36×19 matrix. The columns of **V** (the rows of **V**^*t*^) are an orthonormal basis for those eigenvectors of the recurrent weight matrix **W**_*ŷy*_ that have corresponding eigenvalues equal to 1, and **p** is the projection of **y**_0_ on **V**.

The encoding weight matrix **W**_*zx*_ was a 36×360 matrix (*N*=36 neurons and *M*=360 polar angle samples). The receptive fields (i.e., the rows of the encoding weight matrix **W**_*zx*_) were each designed to be one cycle of a raised cosine:

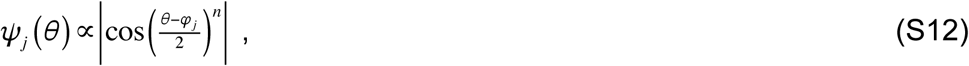

where *θ* is stimulus polar angle and *φ*_*j*_ is the preferred polar angle (i.e., the receptive field center) of the *j*^th^ neuron. The receptive field centers were evenly spaced. The value of the exponent was chosen to be *n* = 18, so that the receptive fields of the 36 neurons spanned a 19-dimensional subspace of polar angles, i.e., the rank of **W**_*zx*_ was 19, equal to the dimensionality of the subspace spanned by **V**. These raised cosine receptive fields have the desirable property that they are shiftable (3, 4), i.e., the response of a neuron with a receptive field of the same shape but shifted to any intermediate polar angle can be computed (exactly) as a weighted sum of the responses of a basis set of 18 of the 36 neurons. The receptive fields evenly tiled the polar angle component of the visual field (i.e., the sum of the squares of the rows of **W**_*zx*_ = 1):

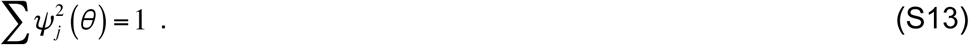

In addition, the encoding weight matrix **W**_*zx*_ was designed so as to project the input stimulus onto the same subspace as that spanned by **V**:

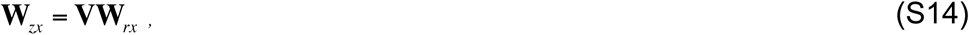

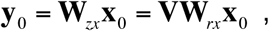

where **x**_0_ was a vector (360×1) corresponding to the input stimulus. The matrix **W**_*rx*_ (19×360) was computed so that the rows of **W**_*zx*_ were the raised cosine functions described above, i.e., **W**_*rx*_ = **V***^t^* **W**_*zx*_.

The same matrix **V** was used to perform the readout. Consequently, the input stimulus could be reconstructed approximately from the steady-state responses:

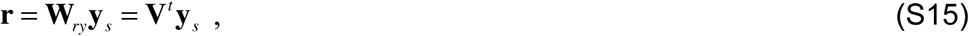

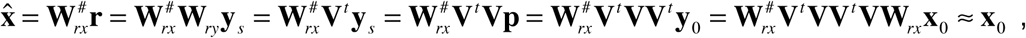

where the last step approximates **x**_0_ because **V** is an orthonormal matrix (i.e., **V**^*t*^**V** = **I**), and # denotes pseudo-inverse. The reconstruction matrix, the pseudo-inverse of **W**_*rx*_, was computed with ridge regression:

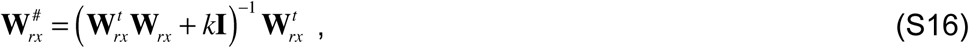

where *k* was a small constant to ensure that the reconstruction was stable with respect to perturbations in **r**.

The input drive was reconstructed exactly from the readout:

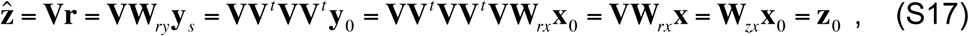

where **z**_0_ is a vector (36×1) corresponding input drive during target presentation.

The steady-state responses (and consequently the readout) were the same even when the encoding weights also included components that were orthogonal to **V**. Specifically, if the encoding weights were **W**_*zx*_ = (**V+V**_*p*_)**W**_*rx*_ such that **V**^*t*^ **V**_*p*_ = 0:

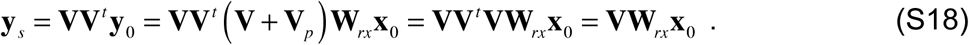

### Oscillatory activity (Fig. 3): readout and reconstruction

The readout for this example circuit was more complicated than that for the circuit that exhibited sustained delay-period activity. The readout depended not only on a weighted sum of the responses but also an estimate of the sign:

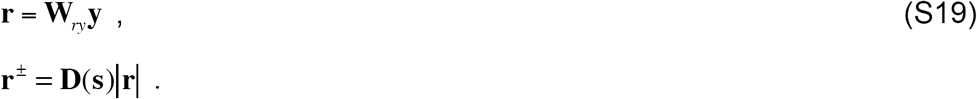

where **r**± is the sign-corrected readout. The matrix **W**_*ry*_ = **V**^*t*^ was a unitary basis for the eigenvectors of the recurrent weight matrix **W**_*ŷy*_ with corresponding eigenvalues that have real parts equal to 1. The vector **s** consisted of ±1 values to correct the sign of the readout, and **D**(**s**) was a diagonal matrix such that each element of the vector **s** was multiplied by the corresponding element of |**W**_*ry*_ **y|**.

The values of **s** were computed from the responses **y**, sampled at two time points. First, the instantaneous frequency of each quadrature pair of neural responses was computed from the real-and imaginary-parts of the responses:

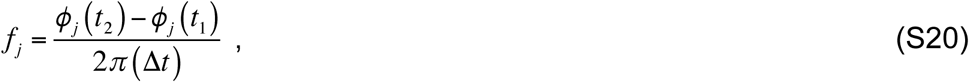

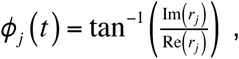

where Δ*t* = *t*_2_ - *t*_1_ was presumed be known, although the values of *t*_1_ and *t*_2_ (i.e., the times at which the responses were sampled) were presumed to be unknown. Second, the elapsed time of the delay period *T* and the corresponding response sign **s** was estimated by minimizing:

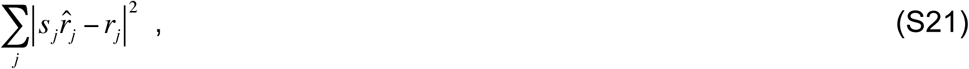

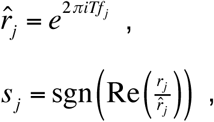

where *r*_*j*_ was the complex-valued response at time *T*, and *f*_*j*_ was the instantaneous frequency (**Eq. S20**). Specifically, we sampled a large number of values of *T* to determine an estimate for the elapsed time that minimized the first line of **Eq. 21**. Given that estimate of *T*, the response sign **s** was then computed using the last two lines of **Eq. 21**. There is a unique solution for **s** when at least two of the oscillation temporal periods have no common multiples. This calculation, as written, is not neurobiologically plausible, but we anticipate that a neural net can approximate the function that transforms from **y** to **s**, or from **y** to **r**^±^.

After correcting the sign of the readout, the input drive and input stimulus were reconstructed using the same procedure as that described above for the sustained delay-period circuit (**Eqs. S15-S17**).

### Sequential activity (Figs. 4 and 5): recurrent weight matrix and reconstruction

The recurrent weight matrix in this example circuit was real-valued but asymmetric. It was computed from a discrete approximation to the derivative of the responses. This was facilitated by using raised cosines for the response fields. The raised cosine receptive fields have the desirable property that they are shiftable (3, 4), i.e., the response of a neuron with a receptive field of the same shape but shifted to any preferred receptive field location can be computed (exactly) as a weighted sum of the responses of a basis set of any 18 of the 36 neurons. First, we computed an upsampling matrix that (exactly) interpolated the responses of the 36 neurons to an arbitrarily large number of responses with arbitrarily closely-spaced preferred locations. Second, we computed a downsampling matrix, as the pseudo-inverse of the upsampling matrix, that subsampled the responses to the receptive field locations of the 36 neurons in the network. Third, we also created a Toeplitz matrix with a convolution kernel that approximated differentiation (5). Fourth, and finally, the recurrent weight matrix depended on the product of these 3 matrices:

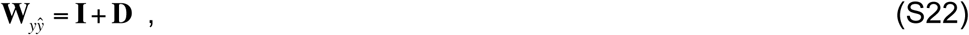

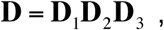

where **I** was the identity matrix, **D**_1_ was the *N*x*M* downsampling matrix, **D**_2_ was the *M*x*M* derivative matrix, **D**_3_ was the *M*x*N* upsampling/interpolation matrix, *N* was equal to the number of neurons (36), *M* was the (arbitrary) upsampling resolution (e.g., *M*=360).

Unlike the preceding examples, it was not possible to reconstruct the input drive from the readout at arbitrary points in time during the delay period. A linear reconstruction (like that used for the preceding examples) generated shifted copies of the input drive, that shifted over time like a traveling wave (**Fig. S4**). Consequently, this circuit was capable of maintaining some (but not all) information about the input during the delay period.

**Figure S4.**
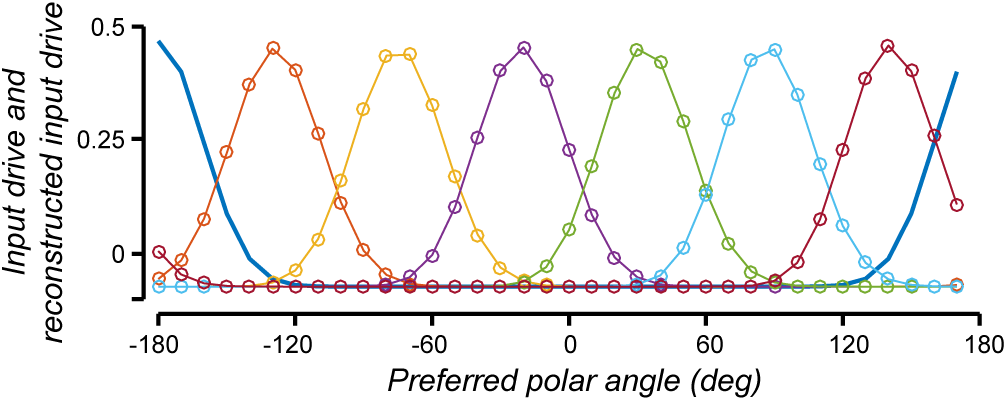
Linear readout and reconstruction for the sequential activity circuit (Fig. 4). Blue curve, input drive (*z*). Other colors, linear readout and reconstruction at evenly spaced time points during the delay period.

### Manipulation (Figs. 6 and 7): recurrent weight *<u>matrices</u>*

There were three recurrent weight matrices in this example circuit, the first for maintaining a representation of the target locations, the second for updating the representation with left-ward eye movements, and the third for updating the representation with rightward eye movements. The first recurrent weight matrix was identical to that in the sustained delay-period activity circuit (i.e., shifted copies of the quadrature-mirror filter kernel, see above). The second recurrent weight matrix was a discrete approximation to the derivatives of the responses (**D** in the 2^nd^ line of **Eq. S22**), and third was the negative derivative matrix (−1 times the matrix defined by **Eq. S22**).

During the period of spatial updating illustrated in **Fig. 6** (i.e., *b* = 0, *a*_1_ = 0, and either *a*_2_ or *a*_*3*_ > 0), **Eq. 4** can be rewritten:

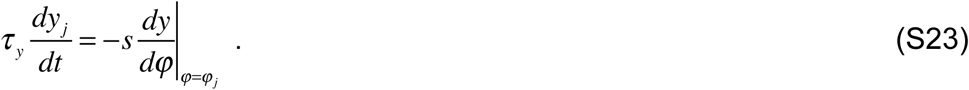

The value of *s* = *a*_*k*_ / (*a*_*k*_ + 1), for *k* equal to either 2 or 3, was the speed of the traveling wave of activity during updating. The variable *φ* was a continuous parameterization of receptive field location (eccentricity), and *y*(*φ*) was a continuous function for every possible receptive field location, interpolated from the *y*_*j*_ samples that represented the neural responses. The derivative d*y*/d*φ*, sampled at *φ* = *φ*_*j*_, was the derivative of the neural responses sampled at the receptive field location of the *j*^th^ neuron. The recurrent weight matrix **D** computed this derivative; multiplying by this weight matrix was equivalent to interpolating the *y*_*j*_ samples to a continuous function, computing the derivative of that function, and then resampling. The duration of spatial updating (i.e., the period of time during which) was proportional to the amplitude of the eye movement:

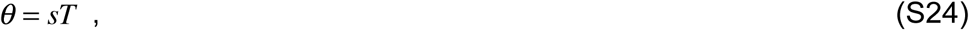

where *θ* was the location of the target stimulus and *T* was the duration of updating. Consequently, the neural responses over time were discrete samples of a continuous function:

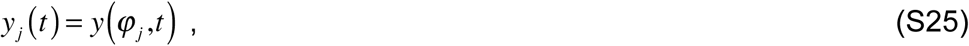

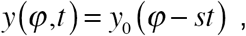

where *y*_0_ was the continuously interpolated neural activity at the point in time (i.e., *t* = 0) just prior to updating.

Likewise, during the period of motor execution illustrated in **Fig. 7** (i.e., *b* = 0, *a*_1_ = 0, and either *a*_2_ or *a*_*3*_ > 0), the responses exhibited oscillating traveling waves. The value of *s* = *a*_*k*_ / (*a*_*k*_ + 1), for *k* equal to either 2 or 3, was the speed of the traveling waves.

### Robustness via normalization (Fig. 8)

ORGaNIC normalization was implemented with a dynamical system comprised of coupled neural integrators:

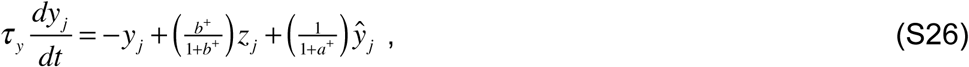

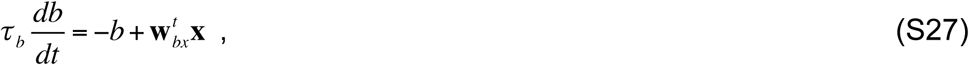

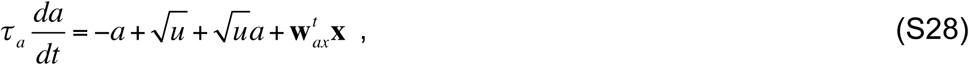

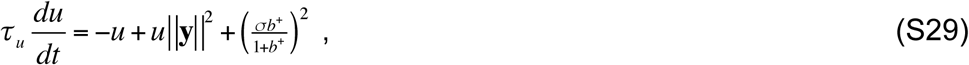

where the norm of *y* is the sum of squares of the real and imaginary parts, summed across neurons:

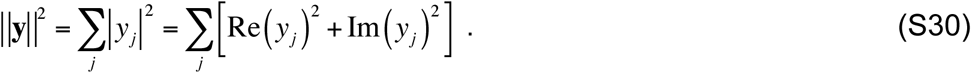

This system of equations is a variation of **Eq. 1**.

As in the preceding examples, all neurons in the circuit shared the same pair of modulators (*a*_*j*_ = *a* and *b*_*j*_ = *b*), i.e., all 36 neurons had the same state at any given point in time. The input modulator *b* depended only on the input, and in particular was set to 1 when a cue was presented indicating the beginning of the trial, i.e., **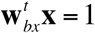** when the cue was presented. The recurrent modulator *a* determined the gain of the recurrent drive. The recurrent modulator *a* depended on the input such that it was set to 1 at the end of the delay period, i.e., **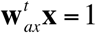** at the end of the trial. The most important difference between this dynamical system and the simpler ORGaNICs expressed by **Eq. 1** is that the recurrent modulator *a* also depended on a particular nonlinear function of of the responses (**Eqs. S28**-**S29**).

The sustained activity and oscillatory activity circuits can achieve a stable state in which the output responses are normalized:

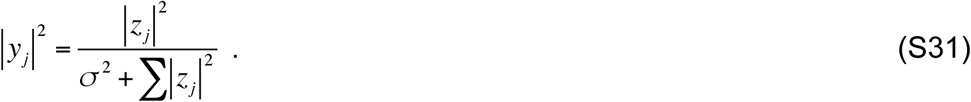

To derive this result, we restrict the analysis to when 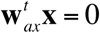 (noting that 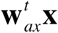 was non-zero only at the end of the trial after the delay period), and when *a* and b are both ≥ 0 (noting that this will generally be the case in the stable state), and we write the stable state for each of **Eqs. S26**, **S28**, and **S29**:

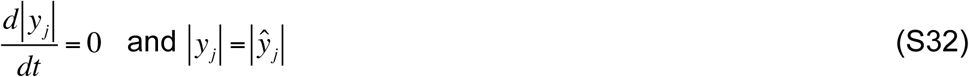

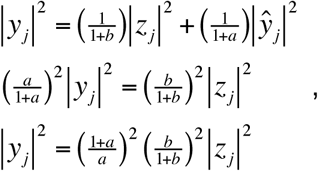

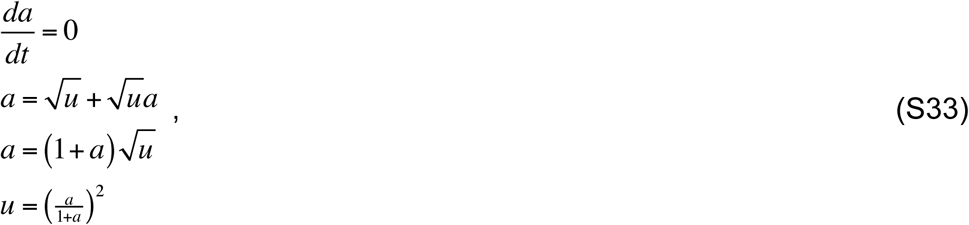

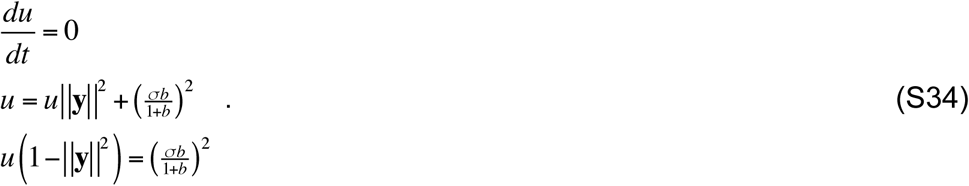

Combining the last line of **Eq. S32** with the last line of **Eq. S33**:

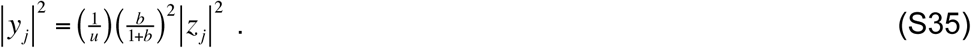

Combining **Eq. S35** with the last line of **Eq. S34**:

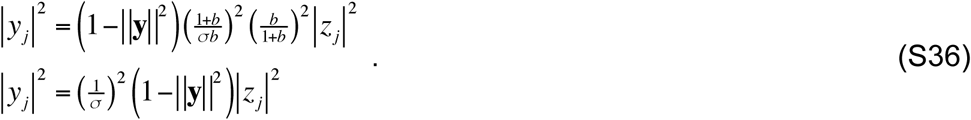

Summing both sides:

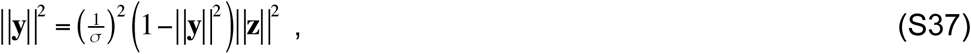

and simplifying:

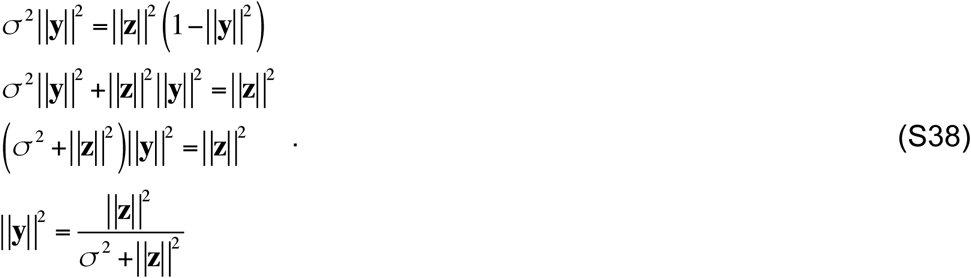

Comparing **Eq. S37** with the last line of **Eq. S38**:

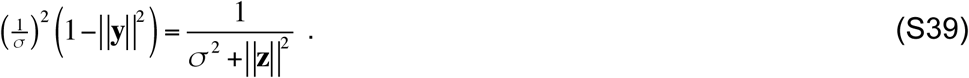

Substituting **Eq. S39** into the last line of **Eq. S36** yields the desired result (**Eq. S31**).

To show that the ratios of the responses equal the ratios of the input drives, we combine **Eq. S31** with the last line of **Eq. S38**:

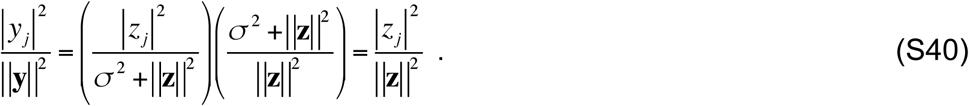

The circuit expressed by **Eqs. S26-S30** is but one example of an ORGaNIC normalization model. There is, in fact, a family of similar dynamical systems models, each of which comprises coupled neural integrators to implement normalization. All of the models in this family achieve the same stable state (**Eq. S31**), but the various different models in this family imply different circuits with different dynamics. For example, some variants including an additional cell type that performs an intermediate step in the computation.

For the sequential activity circuit, the stable state is different:

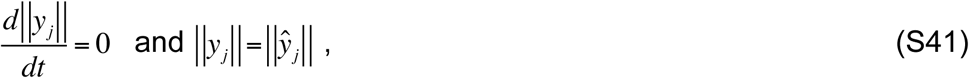

i.e.,

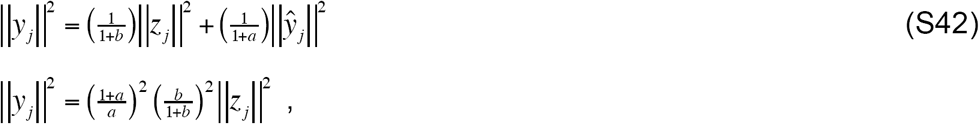

which yields:

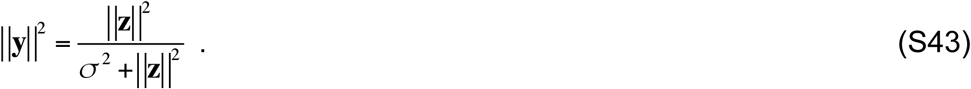

### Variations and extensions

A variant of ORGaNICs is capable of prediction over time (6, 7). Information processing in the brain is dynamic; dynamic and predictive processing is needed to control behavior in sync with or in advance of changes in the environment. Without prediction, behavioral responses to environmental events will always be too late because of the lag or latency in sensory and motor processing. Prediction is a key component of theories of motor control and in explanations of how an organism discounts sensory input caused by its own behavior (8-10). Prediction has also been hypothesized to be essential in sensory and perceptual processing (11-13), and in navigation and the planning of action sequences (14).

ORGaNICs build on Heeger’s Theory of Cortical Function (TCF) (6) that offers a framework for understanding how the brain accomplishes three key functions: (i) inference: perception is nonconvex optimization that combines sensory input with prior expectation; (ii) exploration: inference relies on neural response variability to explore different possible interpretations; (iii) prediction: inference includes making predictions over a hierarchy of timescales. TCF has a single modulator for all of the neurons in each layer whereas ORGaNICs may have a separate pair of modulators, *a*_*j*_ and *b*_*j*_, for each neuron *y*_*j*_. ORGaNICs also have a more general form for the recurrent weight matrix. But TCF includes a feedback drive across the layers of a stacked architecture, in addition to the input drive and recurrent drive. In some states (depending on the values of the modulators), neural responses are dominated by the feedforward drive and TCF is identical to a conventional feedforward model (e.g., deep net), thereby preserving all of the desirable features of those models. In other states, TCF is a generative model that constructs a sensory representation from an abstract representation, like memory recall. In still other states, TCF combines prior expectation with sensory input, explores different possible perceptual interpretations of ambiguous sensory inputs, and predicts forward in time. A variant of ORGaNICs may be stacked (like TCF) to include feedback connections and the capability of a generative model, but with greater flexibility and computational power because of the general form for the recurrent weight matrix, and because there may be a separate pair of modulators for each output neuron.

There is considerable flexibility in the formulation of ORGaNICs, with different variants corresponding to different hypothesized neural circuits (e.g., by a change of variables that might necessitate a different cell type or by including an additional cell type that performs an intermediate step in the computation). For example, we could replace 1/(1+*a*^+^) in **Eq. 1** with 2*a*′/ (1+*a*′), in which 0<*a*′<1. In the original formulation, the activity of the modulator *a*^+^ = 0 during a delay period and non-zero during reset. But in this alternative formulation, the modulator *a*′ = 1 during a delay period and zero during reset. We have implemented and tested many other variants as well; in fact, there is a large family of dynamical systems models, each of which comprises coupled neural integrators, with similar functionality.

The readouts were different for each of the example circuits above. For the sustained activity circuits (**Figs. 2** and **6**), linear readouts reconstructed the input drive (and the target location). For the circuit exhibiting oscillatory dynamics (**Fig. 3**), a (sign-corrected) nonlinear readout was used to reconstruct. For the motor control circuits (**Fig. 5** and **7**), a different linear readout was used to convert spatial patterns of input (premotor) activity to temporal profiles of output (motor control) activity. We did not attempt to reconstruct the input stimulus from the readout, because recovering the input is not the goal for motor control. Likewise, we did not reconstruct the input from the sequential activity circuit (**Fig. 4**). The modulus of the readout enabled the output to be constant over time (i.e., supporting maintenance), but this readout was not capable of reconstructing the input drive. We do not, in general, mean to imply that the brain attempts to reconstruct an input stimulus from the responses. Reconstruction is useful for short-term memory, but not for other cognitive functions. Rather, we hypothesize that the brain relies on a set of canonical (nonlinear) neural computations, repeating them across brain regions so that each stage of computation transforms the information it receives (15-19). The readout is part of this nonlinear transformation from one stage of processing to the next.

There are various formulations for how to compute the modulators. According to **Eq. 2**, the recurrent modulators are a linear sum of the responses **W**_*ay*_ **y**, but the normalization circuit instead computed the recurrent modulators using a nonlinear function of the responses (see above for details). Other variants (particularly for complex-valued responses) compute linear sums of the modulus of the responses, **W**_*ay*_ |**y**|, or linear sums of the various readouts: **W**_*ay*_ **r**, **W**_*ay*_ **r+**, **W**_*ay*_ **r**±. And likewise for the input modulators. According to **Eq. 1**, there is a separate pair modulators *a*_*j*_ and *b*_*j*_ for each neuron, but this need not be the case. Subpopulations of neurons might share some modulators. For example, all 36 neurons shared a single pair of modulators in the various example circuits above. Another option would be to have a number of basis modulators that are shared:

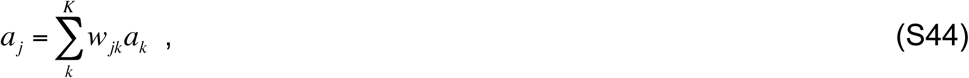

where *a*_k_ are the responses of the basis modulators, *w*_*jk*_ are weights, and the number of basis modulators *K* is less then the number of neurons *N*. And likewise for the input modulators *b*_*j*_. This would dovetail with the idea that some of the modulators are computed via thalamocortical loops (see Discussion), because the number of neurons in a thalamic nucleus is much smaller than the number of neurons in the corresponding cortical area.

The neurons in each of the example circuits in this paper all had the same values for the time constants. That can be generalized, e.g., so that inhibitory neurons have a slower time constant (and consequently delayed responses) compared to excitatory neurons (7). If different neurons in the circuit have different time constants, then the value(s) of the time constant(s), in combination with the eigenvalues of the recurrent weight matrix, determine whether or not there is sustained activity or oscillations, whether the oscillations are stable or decay, and the frequencies of the oscillations.

Optionally, the encoding and readout weights may have components that are orthogonal to the eigenvectors of the recurrent weight matrix (see above for details), and non-zero spontaneous firing rates can be accommodated by adding offsets to each neuron’s input drive (7). For example, if there is a non-zero offset added to the recurrent drive, then the corresponding component of the responses will reflect the elapsed time interval since the beginning of a delay period.

Left open is how to determine the weights in the various weight matrices: the encoding matrix (**W**_*zx*_), the recurrent weight matrix (**W**_*ŷy*_), the readout matrix (**W**_*ry*_), and the modulator weight matrices (**W**_*ax*_, **W**_*bx*_, **W**_*ay*_, **W**_*by*_). Some of the weight matrices (e.g., the recurrent weights) might be pre-specified during development, resulting in stereotypical connectivity within a local circuit. A supervised learning approach would estimate the weights via gradient descent (i.e., back propagation), given target values for the response time-courses (or the readout time-courses), but the brain is unlikely to have access to such target values sampled over time. Another approach would be an unsupervised learning algorithm based on minimizing prediction error over time (6).

### Implications for artificial intelligence and machine learning

1) Go complex. Simple harmonic motion is everywhere. For many AI applications (e.g., speech processing, music processing, analyzing human movement), the dynamics of the input signals may be characterized with damped oscillators, in which the amplitudes, frequencies and phases of the oscillators may change over time. The complex-valued weights and responses in ORGaNICs are well-suited for these kinds of signals. Likewise, we propose using damped-oscillator basis functions as a means for predicting forward in time (6, 7). Traditional LSTMs essentially approximate modulated, oscillatory signals with piecewise constant (or piecewise exponential) functions. There has been relatively little focus on complex-valued recurrent neural networks (20-34), and even less on complex-valued LSTMs (35-39)

2) Stability and vanishing gradients. To ensure stability and to avoid exploding gradients during learning, the recurrent weight matrix should be rescaled so that the eigenvalue with the largest real part is no larger than 1. This rescaling could be added as an extra step during learning after each gradient update. Doing so should help to avoid vanishing gradients by using halfwave rectification instead of a sigmoidal output nonlinearity (38).

3) Normalization. Incorporating normalization can make the computation robust with respect to imperfections in the recurrent weight matrix (**Fig. 8**). Normalization maintains the ratios of the responses (**Eq. 6**), unlike sigmoids or other static output nonlinearities (also called transfer functions) that are typically used in ML systems.

4) Time warping. ORGaNICs offer a means for time warping (**Fig. 7**). Invariance with respect to compression or dilation of temporal signals (e.g., fast vs. slow speech) is a challenge for many AI applications. ML systems typically attempt to circumvent this problem by learning models with every possible tempo. ORGaNICs might be applied to solve this problem much more efficiently, eliminating redundancy and increasing generalization, with less training.

5) Neuromorphic implementation. Given the biophysical (equivalent electrical circuit) implementation of ORGaNICs (**Fig. 8**), it may be possible to design and fabricate analog VLSI ORGaNICs chips. Analog circuitry may be more energy-efficient in comparison to representing and processing information digitally (40, 41). Such an analog electrical-circuit may be configured to download various parameter settings (e.g., the weight matrices), computed separately offline.

